# Leveraging Synthetic Virology for the Rapid Engineering of Vesicular Stomatitis Virus (VSV)

**DOI:** 10.1101/2024.09.26.615165

**Authors:** Chad M. Moles, Rupsa Basu, Peter Weijmarshausen, Brenda Ho, Manal Farhat, Taylor Flaat, Bruce F. Smith

## Abstract

Vesicular stomatitis virus (VSV) is a prototype RNA virus that has been instrumental in advancing our understanding of viral molecular biology and has applications in vaccine development, cancer therapy, antiviral screening, and more. Current VSV genome plasmids for purchase or contract virus services provide limited options for modification, restricted to predefined cloning sites and insert locations. Improved methods and tools to engineer VSV will unlock further insights in long-standing virology questions and new opportunities for innovative therapies. Here, we report the design and construction of a full-length VSV genome. The 11,161 base pair synthetic VSV (synVSV) was assembled from four modularized DNA fragments. Following rescue and titration, phenotypic analysis showed no significant differences between natural and synthetic viruses. To demonstrate the utility of a synthetic virology platform, we then engineered VSV with a foreign glycoprotein, a common use case for studying viral entry and developing anti-virals. To show the freedom-of-design afforded by this platform, we then modified the genome of VSV by rearranging the gene order, switching the positions of VSV-P and VSV-M genes. This work represents a significant technical advance, providing a flexible, cost-efficient platform for the rapid construction of VSV genomes, facilitating the development of innovative therapies.

## 1. Introduction

Vesicular stomatitis virus (VSV) is a negative-sense RNA virus of the Rhabdoviridae family and vesiculovirus genus. The viral genome is non-segmented and approximately 11.1 kilobases (kb) in length, and encodes five genes (N, P, M, G, and L)[1]. VSV is recognized as a livestock pathogen and is well known for its broad tropism and high infectivity. As a prototype virus, it is well studied and boasts a number of favorable features that we have discussed previously in great detail[2]. Highlighting a few of these, VSV has a relatively simple genome, it readily accepts foreign glycoproteins, and it is known for fast replication kinetics[3]. With replication cycles of 4–6 h, VSV and variants can be produced at high titers with relative ease[4, 5]. Taking these into consideration, it is not surprising that VSV has been instrumental in advancing our understanding of viral molecular biology and has become an essential tool in the research and development of new vaccines and therapies.

This special issue delves into novel insights regarding VSV and explores how this virus can be harnessed as a tool to combat infectious diseases, cancer, and other maladies. VSV has a proven track record in vaccine development. ERVEBO, approved by the FDA in 2019, is a recombinant VSV expressing an Ebola virus glycoprotein[6]. Additional studies have explored VSV-based vaccine vectors for HIV[7-9], Chikungunya[10, 11], and SARS-CoV-2[12], among others[1, 13]. Another article in this special issue developed a VSV-based Marburg vaccine[14]. VSV pseudotypes have been used to study tropism, cell entry, and even to develop antivirals against these corresponding glycoproteins[14, 15]. Of note, pseudotyping is when the natural glycoprotein sequence of a virus is deleted from the genome, and then they are packaged with a foreign glycoprotein[16, 17]. Beyond vaccines and antivirals, VSV encoding an interferon beta transgene (VSV-IFNB) is currently in clinical trials to treat solid tumors[18]. Despite the diverse applications and the wide range of VSV variants, they share a common origin: their viral genomes were engineered, constructed, and subsequently converted into virus particles, a process known as virus rescue or recovery. However, virus engineering is inherently difficult, and further limited by available materials and tools.

Current methods and technologies for engineering viruses are limited by long time periods, the need for significant scientific expertise, and financial resources[19-21]. Traditional manipulation of viral genomes requires homologous recombination in plasmids, along with subsequent selection and validation, which typically restricts construction to sequential steps[20, 22, 23]. Restriction enzyme based cloning methods are limited to locations of restriction sites. New ones can be generated, by PCR, site directed mutagenesis and other methods, but this adds time and validation steps. Incomplete digestion, DNA methylation, compatibility of restriction sites, and difficulties with multiple digests are some of the factors contributing to the limitations and issues that arise with this approach[24-26]. Advanced engineering approaches rely on DNA editing, primarily CRISPR-Cas systems, which have a number of concerns and limitations, including: varying guide specificity, unintended mutagenesis, off-target activity, low efficiency when working with multiple targets, and the need for consecutive rounds of editing when incorporating multiple changes in genetic constructs[27-33]. Taken together, there is a significant unmet need for new virus engineering technologies and drug development strategies.

Synthetic virology is the engineering of virology: the rational design and construction of new viruses and systems inspired by viruses with functions not found in nature. The chemical synthesis of infectious poliovirus in 2002 by Eckard Wimmer demonstrated the feasibility and utility of whole-genome synthesis and established the field of synthetic virology[34]. This landmark study paved the way for the de novo synthesis and assembly of other viral genomes, including bacteriophage ΦX174, the 1918 Spanish influenza pandemic virus, tobacco mosaic virus (TMV), horsepox virus, Herpes simplex virus type 1 (HSV-1)[35], among others[27, 36-49]. Scientists are using synthetic virology for biomedical research (e.g., functional studies, archaevirology)[27], improving viral vectors for gene therapy[50, 51], reprogramming bacteriophages as antibiotics and therapeutics[52, 53], vaccines[54, 55], and more. For example, Thao et al. reconstructed synthetic SARS-CoV-2 for viral research and vaccine development[56], and Huang et al. assembled a synthetic oncolytic adenovirus with genetic circuitry for the treatment of hepatocellular carcinoma[57]. VSV’s adaptability for use in vaccines and cancer therapies necessitates specific modifications to tailor it for distinct applications. To this end, there is an unrealized opportunity to leverage synthetic virology to enable rapid engineering of VSV genomes.

In this work, we introduce a methodological framework that facilitates the rapid design, construction, and testing of synthetic viruses (Figure 1). This paper highlights a significant technical advancement by merging computer-aided biological design tools with comprehensive whole-genome synthesis and assembly techniques to generate viable, replication-competent viral particles. This novel approach is not only scalable and cost-effective but also maintains logical consistency throughout the process.

**Figure 1.**
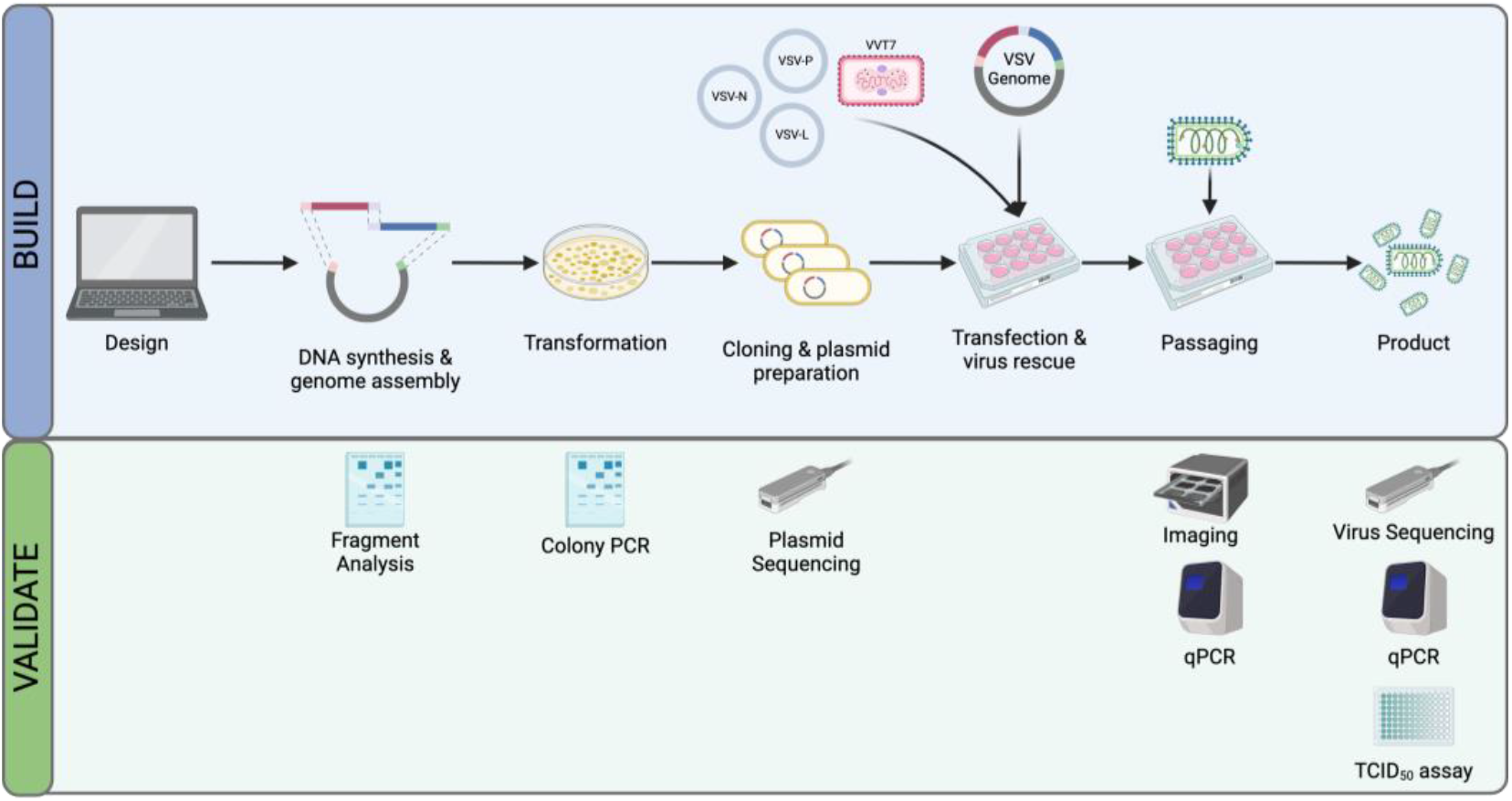
Overview of the VSV engineering platform. Internal bioCAD, or other DNA visualization software tools (e.g., SnapGene), are used to design complete VSV genome. The design spec if broken down into fragments with modular adapters, may be reused in future designs. Fragments are synthesized, and DNA is then assembled into complete plasmid. This is transformed into bacteria, propagated, and purified. Viral genome plasmids are validated by colony PCR and whole plasmid sequencing throughout. These VSV genome plasmids are transfected in vaccinia virus-T7 infected 293T cells along with support plasmids expressing VSV N, P and L proteins. Virus is recovered, validated, and expanded as needed to increase titers.

## 2. Materials and Methods

### 2.1. Viruses

Vesicular stomatitis virus (VSV), Indiana strain was obtained as a complete plasmid from Kerafast (Kerafast, Shirley, MA, USA, Cat #EH1002) and rescued by traditional reverse genetics per suggested protocol from depositor (the laboratory of Michael Whitt, Memphis, TN, USA). This was used as the reference virus for VSV wildtype. Vaccinia virus, western reserve strain encoding a bacteriophage T7 RNA polymerase transgene, also known as VVT7 or vTF7-3, was purchased from Imanis Life Sciences (Cat #REA006) for rescue of VSV from plasmid DNA by reverse genetics.

### 2.2. Plasmids

VSV (Kerafast, Shirley, MA, USA, Cat #EH1002), a full-length Indiana strain VSV plasmid, was used to rescue wild-type virus. This was used as the reference virus for VSV wildtype and considered the natural isolate or variant in this article. Additionally, helper or support plasmids, designated as pBS-N, pBS-P, pBS-G, and pBS-L (names correspond to each VSV gene, inserted in a BlueScript SK plus, or pBS-SK-ΦT, cloning vector) were obtained from Kerafast. Expression of these VSV genes is dependent on the presence of T7 polymerase, which is traditionally supplied by vaccinia virus described in the previous section.

### 2.3. Computer-Aided Design

The first step of the design process was obtaining a virus reference sequence for vesicular stomatitis virus (VSV), Indiana serotype; This was provided by Kerafast, (Kerafast, Shirley, MA, USA) plasmid service provider. Whole plasmid sequencing was performed by Plasmidsaurus to determine VSV wildtype plasmid, as confirmation of the provided reference sequence (Figure 2). Next, BioCAD software (Humane Genomics developed) for virus engineering, and the VSV reference sequence was deconstructed digitally into potential fragments for subsequent DNA synthesis and assembly.

**Figure 2.**
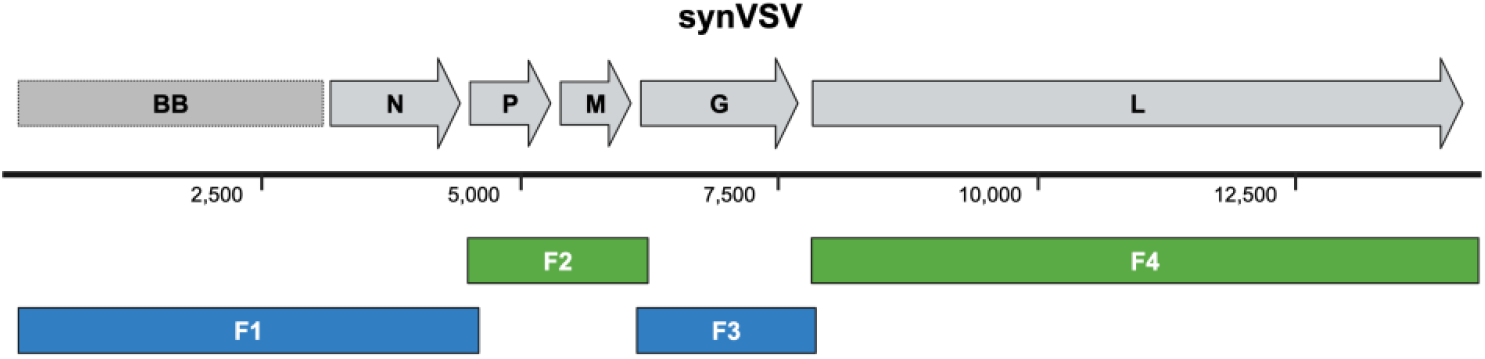
Genome schematic of vesicular stomatitis virus (VSV), Indiana serotype reference genome is depicted on top, along with a reference bar indicating genome size (number is in reference to base pairs). During the design step, this genomic sequence is then deconstructed into DNA fragments that are then submitted for synthesis. Depicted in blue, these fragments have 30 base pair overlap sequences that enable subsequent assembly into a complete, physical DNA genome. This final circularized synVSV genome plasmid is depicted in blue at the bottom.

Fragment start and stop sites were determined based on three main criteria: the location (at the beginning or end of coding sequences), the size (based on the number of base pairs), and the compatibility with commercial DNA synthesis platforms. To enable one-pot homologous assembly, 30 base pair overlap sequences were designed at the terminal 5′ and 3′ ends of the fragments. These unique sequences ensured precise assembly with no byproducts or off-target recombination, controlling fragment directionality during assembly to obtain the desired product. Importantly, the rational design of these fragments allows for reuse of long DNA sequences or constant viral elements that do not change across different designs, reducing the overall cost of DNA synthesis over time.

Apart from the VSV genome and the overlapping fragments used for assembly, the design incorporated the BlueScript SK plus (pBS-SK-ΦT) cloning vector backbone, similar to the commercially available plasmid from Kerafast (Kerafast, Shirley, MA, USA).

Several restrictions and challenges are associated with DNA synthesis services, including limits on sequences with repeats longer than 10 base pairs, fragments with melting temperatures above 60 °C, GC content below 25% or above 65%, and variations in local GC content (with differences greater than 52% between the highest and lowest 50 base pair regions).

For the design of VSV with the Sindbis virus glycoprotein (VSV-SIN), the envelope sequence from Sindbis virus strain TE3′2J (AR339) was used. Overlap sequences matching the VSV P-M fragment (F2) were incorporated at the 5′ end of the glycoprotein fragment (sequence: *aaactaacagagatcgatctgtttacgcgt*), and a sequence matching the VSV-L fragment (F4) was included at the 3′ end (sequence: *aacagcaatcatggaagtccacgattttga*).

For the design of VSV with gene rearrangement (VSV-M2P3), the overlap sequences matching the VSV N fragment (F1) were used at the 5′ end of the MP fragment (sequence: *tgacaaatgaccctataattctcagatcac*), and a sequence matching the VSV-G fragment (F3) was included at the 3′ end (sequence: *aaactaacagagatcgatctgtttacgcgt*).

### 2.4. DNA Synthesis and Genome Assembly

The FASTA files of our designed fragments were uploaded to Twist Bioscience for DNA synthesis. After receiving the resulting linear DNA fragments (0.3–7 kb), they were quantified by UV-vis spectrophotometry using a Nanodrop (ThermoFisher, Waltham, MA, USA). Concentration was determined by measuring absorbance at 260 nm (A260, nucleic acid concentration), and A260/A280 ratio was used to evaluate quality. Next the DNA fragments were run on Invitrogen precast 0.8% agarose NGS gels for visual confirmation of size.

Four DNA fragments, representing the VSV antigenome and plasmid backbone of pBS, were combined and assembled using NEBuilder HiFi DNA Assembly Master Mix (New England Biolabs, Ipswich, MA, USA). This reaction consisted of DNA (0.05 pmol per fragment), 10 µL of master mix, and nuclease free water (NFW) if needed, to a final volume of 20 µL. This reaction was set up on ice. Following preparation, reactions were incubated on a pre-heated thermocycler at 50 °C for 60 min, per the manufacturer’s recommendation. Afterwards, NEB stable *E. coli* competent cells (New England Biolabs, Ipswich, MA, USA) were chemically transformed with 20 µL of assembled DNA product, streaked on agar plates supplemented with antibiotics, and evaluated the next day for colony growth.

### 2.5. Colony PCR & Gel Electrophoresis

Colony PCR was performed to evaluate the presence of the VSV-G gene using 2x Q5 hot start high fidelity master mix (Invitrogen, Waltham, MA, USA) as per manufacturer’s instructions. In short, 10 colonies were picked after overnight incubation of the transformation plates at 30 °C. The colonies were inoculated in 10 µL of nuclease free water. Next, 1 µL of the inoculum was used as template for the colony PCR reaction. For the colony PCR set up in the PCR tubes, 12.5 µL of Invitrogen 2x Q5 hot start high fidelity master mix was mixed along with 10 µM of VSV-G specific forward (5′-ggacctacgcgtatgaagtgccttttgtacttagcc-3′) and reverse (5′-aggcccgctagcctttccaagtcggttcatctctat-3′) primers along with 1 µL of template. The remaining volume was adjusted to 25 µL by addition of nuclease free RT-PCR grade water (Invitrogen, Waltham, MA, USA). Then 20 µL of PCR amplicons were run on 0.8% precast ready to use agarose NGS e-gel (Invitrogen, Waltham, MA, USA) alongside a Thermo Fisher Generuler 1 Kb plus molecular size DNA ladder to identify the positive colonies of ~1.5 kb amplifying the target gene (VSV-G). The gel was run for 26 min, and images were captured automatically five times throughout.

### 2.6. Plasmid Miniprep

The remaining 9 µL of the corresponding positive bacterial inoculums in nuclease free water (from the initial step of colony PCR) were used to inoculate 5 mL ampicillin containing LB media, overnight cultures were grown at 30 °C and plasmid minipreps were performed the following day. Plasmid minipreps of the overnight bacterial cultures corresponding to the VSV genome containing colonies were performed using Qiagen QIAPrep spin miniprep kit (Qiagen LLC, Germantown, MD, USA) as follows. 5 mL overnight bacterial cultures were transferred to 1.5 mL microcentrifuge tubes and centrifuged at 6800× *g* in a high-speed centrifuge. The pellets were resuspended, lysed, neutralized and washed with the kit’s buffers following the Quick-start protocol provided in the handbook. The final plasmids DNA were eluted in 50 µL volume in sterile nuclease free microcentrifuge tubes. The quality & concentration of the purified plasmid minipreps were determined by UV-vis spectrophotometry using a Nanodrop 260/280 ratio values (ThermoFisher, Waltham, MA, USA). Whole plasmid sequencing of viral genome plasmids was performed by Plasmidsaurus using Oxford Nanopore Technology with custom analysis and annotation.

### 2.7. Cells

Four cell lines were used in the process and purchased from ATCC: 293T (CRL-3216), HepG2 (HB-8065), and Hep3B (HB-8064). All cell lines were maintained per the supplier’s recommendations. The 293T cells were cultured in Dulbecco’s Modified Eagle Medium (DMEM) containing L-glutamine, phenol red and high glucose supplemented with 10% fetal bovine serum, 1% Penicillin-Streptomycin antibiotic (2500 units) and 1% antimycotic antibiotic. The HepG2 and Hep3B cells were maintained & cultured in Eagle’s Minimum Essential Medium (EMEM) containing L-glutamine, phenol red and high glucose supplemented with 10% fetal bovine serum, 1% Penicillin-Streptomycin antibiotic (2500 units) and 1% antimycotic antibiotic. HepaRG (NSHPRG) were obtained from Lonza, and maintained in HepaRG growth medium per manufacturer’s recommendation. All cells were maintained in an incubator set at 5% CO_2_, 37 °C. Routine mycoplasma testing was conducted (ATCC, MY01050).

### 2.8. Virus Rescue

Following the design, construction, and validation of VSV genome plasmids, functional virus particles were generated and recovered using traditional reverse genetics methods of vaccinia virus-mediated delivery of bacterial T7 polymerase. In brief, 293T were seeded in 12-well plates (2 × 10^5^ cells per well), incubated for 18–24 h, and then infected with vaccinia virus with a MOI of 0.5 TCID_50_ (293T). After 45 min of inoculation at 37 °C, virus-containing media was removed. Cells were then transfected with desired VSV genome plasmid (1 µg), pBS-N (0.5 µg), pBS-P (0.4 µg), pBS-G (1 µg), pBS-L (0.2 µg) using Lipofectamine 3000 (Invitrogen) per manufacturer’s recommendations. After 48 h, cells were rounded and detaching due to a combination of vaccinia virus and VSV mediated cytopathic effects (CPE).

Viral supernatant was harvested, centrifuged for 10 min at 450× *g*, and then the supernatant was filtered with a 0.22 µm filter to remove vaccinia virus prior to amplification. Virus was subsequently passaged in a 12-well plate of 293T (2 × 10^5^ cells per well) that were seeded 18–24 h prior to infection. Signs of cytopathic effects and rounding at 12–72 h post-infection were indicative of successful rescue. To this end, viral supernatant was harvested, centrifuged for 10 min at 450× *g* to remove cell debris, and then the supernatant was aliquoted and stored at −80 °C. These viral stocks were used for further confirmation: RT-qPCR using VSV-M primers for initial confirmation of VSV-specific genome presence, nanopore sequencing for full genome sequence validation. After passing QA/QC inspection, stock aliquots were then used for virus characterization: RT-qPCR using VSV-M primers to quantify genome copies, TCID_50_ assay for titration and quantification of infectious virus. More details on these assays are provided in the following sections.

### 2.9. Quantitative Real-Time Reverse Transcription PCR (RT-qPCR)

RNA was extracted from cell culture supernatant using QIAamp^®^ DSP Viral RNA Mini Kit (QIAGEN LLC, Germantown, MD, USA, Cat No. 61904). This extracted RNA was used for confirmation of recovery and quantification of virus genome copies, which is described below.

RNA was measured using Nanodrop for concentration and quality. First strand cDNA synthesis was then carried out in a 20 µL reaction containing 50 µM Random Hexamer Primer (Thermo Scientific, Cat No. SO142), 10 mM dNTP, 5X Induro RT Reaction Buffer, 200 U/µL Induro^®^ Reverse Transcriptase (NEB, Ipswich, MA, USA, Cat No. M0681), 20 U/µL SUPERase-In Inhibitor, and NFW. A maximum of 40 ng or 10 µL RNA was used per reaction. The following cycling parameters were used: 2 min at 25 °C, 20 min at 55 °C, followed by 1 min at 95 °C. cDNA was then measured for VSV-M using SYBR Green qPCR on a QuantStudio 6 Flex instrument (ThermoFisher). The 20 µL reaction contained 1 µL cDNA, 0.5 µL Yellow Sample Buffer, 10 µL, PowerTrack SYBR Green Master Mix, 200 nM VSV-M forward and reverse primer, and NFW. The following sequences were used: VSV-M forward primer 5′-TGGAGTTGACGAGATGGACAC-3′; VSV-M reverse primer 5′-TTTCCCTGCCATTCCGATGT-3′. The following cycling parameters were used: one cycle of enzyme activation (2 min at 95 °C), followed by 40 cycles of Denaturation (15 s at 95 °C) and Annealing/Extension (1 min at 55 °C). A standard curve of synthetic DNA fragments representing 1 × 10^3^ to 1 × 10^8^ VSV-M gene copies (ten-fold dilutions) was generated and used. Applied Biosystems instrument software was used to calculate genome copy number.

### 2.10. *TCID*_*50*_ Assay

Tissue Culture Infectious Dose 50 (TCID_50_) assay was used to quantify functional, infectious virus and determine VSV titer for known and unknown samples. In brief, 293T cells were seeded in 96-well plates at a density of 10^4^ cells per well and incubated overnight at 37 °C with 5% CO_2_. Virus was thawed at room temperature and stored on ice for assay preparations. Serial tenfold dilutions of the VSV stock aliquot were prepared in complete media. Each dilution was added to the wells in triplicate, and the plates were incubated for 72 h. Cytopathic effect (CPE) was then observed and recorded under a microscope. Immunocytochemistry (ICC) staining was performed at the endpoint using anti-VSV (Imanis Life Sciences, Rochester, MN, USA) to confirm visual observation; More details on ICC are provided in subsequent sections. The TCID_50_ value was calculated using the Spearman-Karber improved method, determining the dilution at which 50% of the wells exhibited CPE. Of note, each run included a negative control or mock infection, and experiments were performed in triplicate to ensure reproducibility and accuracy.

### 2.11. Immunocytochemistry (ICC)

To confirm the presence of VSV, virus was detected in cell assays using immunocytochemistry (ICC). In brief, cells were fixed with 4% formaldehyde (MilliporeSigma, Burlington, MA, USA, Cat#47673) solution for 15 min 25 °C, permeabilized with cold 70% ethanol (ThermoFisher) solution for 15 min. Cells were washed one time with a wash buffer (PBS + 1% BSA [10 g for 1 L] + 1 mM EDTA [292.2 mg for 1 L]) between these steps. Cells were then incubated with a blocking buffer (PBS + 1% BSA + 1 mM EDTA) + 2% FBS) for 45 min at room temperature. Cells were then washed once more before addition of antibodies. Cells were incubated with primary antibody rabbit anti-VSV (Imanis Life Sciences; Cat# REA005) for 60 min at room temperature. Cells were washed 3 times with wash buffer. Then, cells were incubated with secondary antibody Alexa Fluor 488 goat anti-rabbit IgG (H + L) (ThermoFisher; Cat# A11034) for 45 min in the dark at room temperature. Three washes were then carried out before imaging.

### 2.12. Live-Cell Imaging and Analysis

The Incucyte Live-Cell Analysis System (Sartorius) was used for real-time imaging and analysis of cell label-free evaluation of cell proliferation (confluence percentage) and health over time. For mock and virus-infected wells of cells (e.g., TCID_50_ assay), tissue culture plates were scanned using the Incucyte ZOOM imaging system (Version 6.2.9200.0) with a 10× objective lens. The Incucyte ZOOM software (Version 6.2.9200.0) was set up to capture 16 images per well for 6-well plates, 9 images per well for 12-well plates, and 3 images per well for 96-well plates. HD phase and GFP images were captured every 2–6 h on an automated schedule. Afterwards, relevant metrics (e.g., confluence % or cell area) were analyzed per image and averaged per well, and then exported to Google Sheets and GraphPad Prism 9 (Version 9.0, GraphPad Software, San Diego, CA, USA) for further analysis of technical and experimental replicates. Representative images from relevant timepoints were also exported as .PNG files.

### 2.13. Statistical Analysis

The qPCR data are presented as the mean ± SD. Student *t*-tests were performed for comparison. GraphPad Prism 9 (GraphPad Software, San Diego, CA, USA) was used for statistical analyses.

## 3. Results

### 3.1. De Novo Design and Assembly of synVSV Genome

A commercially available VSV wildtype plasmid from Kerafast was used as the template and reference sequence for designing a complete VSV genome that could be assembled de novo. The genome was divided into four DNA fragments, labeled F1 through F4 (Figure 2), which were synthesized by Twist Biosciences. Fragment 1 (F1) spanned the plasmid backbone (pBS-SK-ϕT), the T7 promoter, and the VSV-N coding sequence, totaling 4481 bp. Fragment 2 (F2) was 1751 bp long and included a 30 bp overlap with F1, the VSV-P protein sequence and its transcription unit, the VSV-M protein sequence and its transcription unit, and a 30 bp overlap with F3 located in the 5′ UTR of VSV-G. Fragment 3 (F3) contained the VSV-G protein and its transcription unit. Fragment 4 (F4) was a 6439 bp fragment encompassing the VSV-L gene. The 3′ end of F4 contained a 30 bp overlap with the 5′ end of F1, allowing for the circularization of the fragments upon assembly. Detailed information on each DNA fragment, including sequence and length, is provided in Supplementary Table S1.

After design and synthesis, the fragments were assembled using HiFi DNA assembly, with optimal quantities for each fragment calculated using the NEBiocalculator. Specifically, 0.05 pmol of each fragment was used for the assembly reaction, followed by chemical transformation. Figure 3A shows the efficiency of the reaction, evidenced by the dense lawn of transformed bacterial colonies on the experimental agar plates (pUC19 served as a positive control, though images are not shown). Colonies were then picked and screened for the presence of VSV-G using colony PCR, which identified a distinct 1500 bp band in all ten colonies examined (Figure 3B). This confirmed assembly and demonstrated that all the clones produced had the correct fragments included.

**Figure 3.**
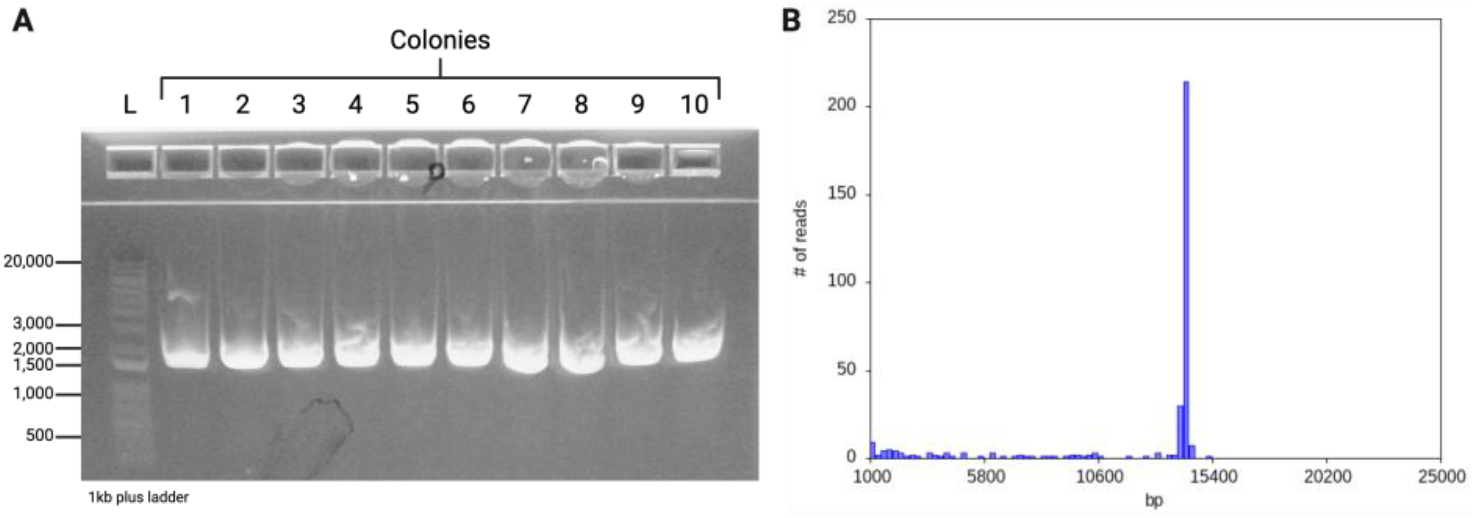
Confirmation of genome assembly. (**A**) Colony PCR was conducted and 10 independent clones were evaluated for the presence of VSV-G. As shown here, the gel is showing positive confirmation of VSV-G for 10/10 clones, indicating 100% efficiency. Colony PCR indicates the positive presence of an amplified target, but for even greater validation, whole plasmid sequencing was performed by Plasmidsaurus. Viral genomes were evaluated for identity and quality. All clones were sequence verified, and no mutations were present. (**B**) Raw read length histogram generated from plasmid sequencing run. Here, a randomly selected clone demonstrated a single peak at the correct length (base pairs), indicating high quality DNA.

Subsequent culture and plasmid preparation were followed by whole plasmid sequencing conducted by Plasmidsaurus. The consensus sequence matched the expected VSV reference sequence, further confirming 100% accuracy in synthetically recreating this virus.

### 3.2. Rescue and Characterization of Synthetic VSV (synVSV)

Functional virus particles were generated using reverse genetics by infecting 293T cells with vaccinia virus to deliver T7 polymerase, followed by lipofectamine-based transfection with the purified synVSV genome and required support plasmids. After 48 h, harvested viral supernatant was purified to remove residual helper vaccinia virus, which was then used for infection of 293T cells (passage 1). Here, we observed 75% of transfected wells were successful in virus recovery, and synVSV demonstrated complete lysis in these wells in less than 24 h. Viral supernatant was harvested from these wells and titered for downstream assays.

Characterization of synVSV was performed to compare its functionality with that of the wild-type virus and to identify any potential variations. To this end, we conducted a standard TCID_50_ assay on production cells (293T) to determine the infectious viral titer. During the assay, we observed the rapid onset and progression of cytopathic effects (CPE), characteristic of VSV. After 72 h, immunocytochemistry (ICC) was performed using an anti-VSV primary antibody to confirm that the observed cell rounding and death were attributable to VSV. It is well-known, and we have also observed, that 293T cells easily detach from plastic surfaces even with light agitation, which can artificially inflate viral titers in a TCID_50_ assay. Therefore, validation by ICC is crucial for ensuring greater accuracy in quantification. However, in our experiment, we encountered issues where wells were either completely lysed, leaving no cells to stain for VSV, or entirely confluent and negative for viral presence. As a result, ICC did not enhance scoring accuracy in this instance. Nonetheless, the observation of complete lysis or its absence allowed us to score the assay, resulting in a titer of 1.5 × 10^10^ TCID_50_/mL (293T), with a 95% confidence interval ranging from 4.3 × 10^9^ to 5.1 × 10^10^.

Viral quantification by TCID_50_ assay was also performed on HepG2 cells, a more adherent cell line. Similarly, staining with an anti-VSV antibody was conducted after 72 h. As expected, the lowest virus dilution (positive in 1 of 3 wells) showed the onset and progression of virus-induced CPE and lysis, confirmed by ICC. The calculated TCID_50_ here was 6.8 × 10^9^ (HepG2), with a 95% confidence interval ranging from 3.2 × 10^9^ to 1.5 × 10^10^.

However, a number of healthy, uninfected cells remained in the wells. Representative images are shown in Figure 4A–D.

**Figure 4.**
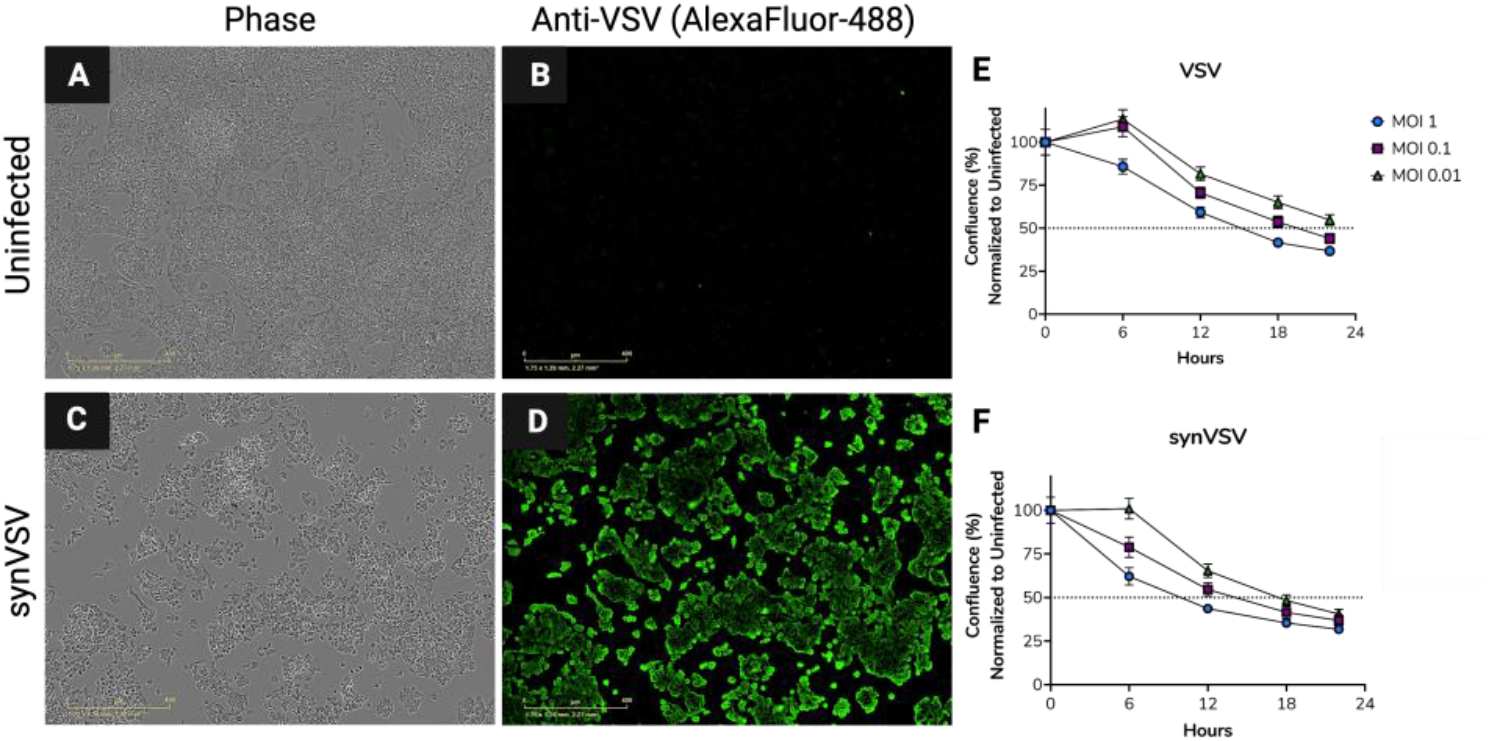
Quantification and characterization of synVSV. (**A**–**D**) Following TCID_50_ assay on HepG2 cells, ICC was performed using anti-VSV antibody to further validate virus presence and virus-dependent cytotoxicity. (**A**,**B**) a random uninfected well from the TCID_50_ assay and ICC was selected. As seen here, cells are confluent and no staining of VSV proteins (indicating VSV presence) was observed. (**C**,**D**) Cells are rounded and showing cytopathic effects at the lowest positive dilution in the assay. Corresponding images taken with green fluorescence channel for AlexaFluor-488 further confirms these negative cell effects are due to the presence of synVSV. (**E**) VSV wildtype (natural/parental) and (**F**) synVSV (synthetic) were infected as multiple dilutions, and viral kinetics were compared to each other. Here, we see a faster onset of CPE and lysis in the synVSV infected wells compared to each corresponding matched MOI in the natural counterpart.

To compare replication dynamics of synVSV and natural counterpart, cell health and proliferation assays were carried out on Hep3B cells with each virus at multiple MOIs (serial ten-fold dilutions starting at MOI = 1). Real-time imaging was conducted using the IncuCyte system, and cell confluence (%) was assessed at 6-h intervals up to 24 h post-infection. Confluence was normalized to uninfected wells, which represent healthy cells and normal proliferation. Consistent with observed cytopathic effects (CPE), consisting of rounded cells and lysis, confluence decreased in a MOI-dependent manner for both viruses (Figure 4E,F). Interestingly, synVSV exhibited faster onset of virus-induced CPE and decline in cell health than VSV wildtype (natural counterpart). This was observed across all MOIs matched with each other.

This phenomenon may be occurring for several reasons, but two stand out. First, when quantifying each virus, ten-fold serial dilutions were used for titration. It’s possible that the synthetic VSV titer is slightly higher than the wildtype, though less than a ten-fold difference, which would result in synVSV having a marginally higher MOI during infection. Second, there may be inherent benefits to using DNA synthesis and genome assembly compared to traditional methods. synVSV could possess optimized properties or kinetics due to subtle differences, such as increased genome stability, fewer mutations, or reduced population variation at the outset. Synthetic DNA is sequence-validated at the start, whereas PCR or amplification-based approaches can introduce mutations and errors. These factors could enhance viral infectivity and replication by providing a higher-quality, more homogenous viral population. Importantly, this detail might not be captured in titration, as plates are read 72 h post-infection, meaning that a slower-replicating virus could still catch up and display the same titer at the endpoint. In this study, viral kinetics were observed up to 24 h, where both synthetic and natural viruses showed a similar downward trend in the confluence graph, though staggered by a few hours.

In summary, the de novo assembled synVSV was functional, with viral yields from production falling within the expected titer range. Robust infection and replication were observed in both 293T, HepG2, and Hep3B cell lines. In the latter, synVSV exhibited superior kinetics compared to VSV wildtype, the natural counterpart. Of note, Xiao et al.[35] similarly reported enhanced properties of synthetic HSV compared to parental viruses recently, but further investigation is warranted for why this is the case.

### 3.3. Evaluate VSV with Foreign Glycoprotein Using the Synthetic Virology Platform

After demonstrating proof-of-concept with synVSV. We wanted to validate the platform further and how it may be used to rapidly construct new VSVs. Given the versatility and power of VSV as a vector for vaccines and medical countermeasures and popularity in labs, we decided to first validate the platform by designing and constructing a VSV with a foreign glycoprotein. After some consideration, we decided to pursue the design and construction of a replication-competent VSV with Sindbis envelope glycoprotein here. Previous groups have evaluated VSV pseudotyped with Sindbis glycoprotein and HER2 targeted ligand[58], as well as replication competent VSV with alphavirus glycoproteins (CHIKV)[59]. However, to the extent of our knowledge, a replication-competent VSV Sindbis chimeric virus has not been evaluated to date.

To design and construct synVSV-ΔG SIN-E3/E2/6K/E1, the overlap sequences on the ends of VSV-G were maintained and a DNA fragment synthesized that contained the Sindbis envelope or structural polypeptide sequence (AR339 strain was used as a reference sequence) (Fig 5). HiFi assembly was performed under the same conditions as synVSV noted prior, QA/QC was conducted and transformants verified, and virus was recovered. Following sequence confirmation of the rescued virus, synVSV-ΔG SIN-E3/E2/6K/E1 virus was evaluated on production cells as well as liver cancer cells. Of note, Sindbis virus has been reported to target the laminin receptor (LAMR), which is highly expressed in hepatocellular carcinoma.

**Figure 5.**
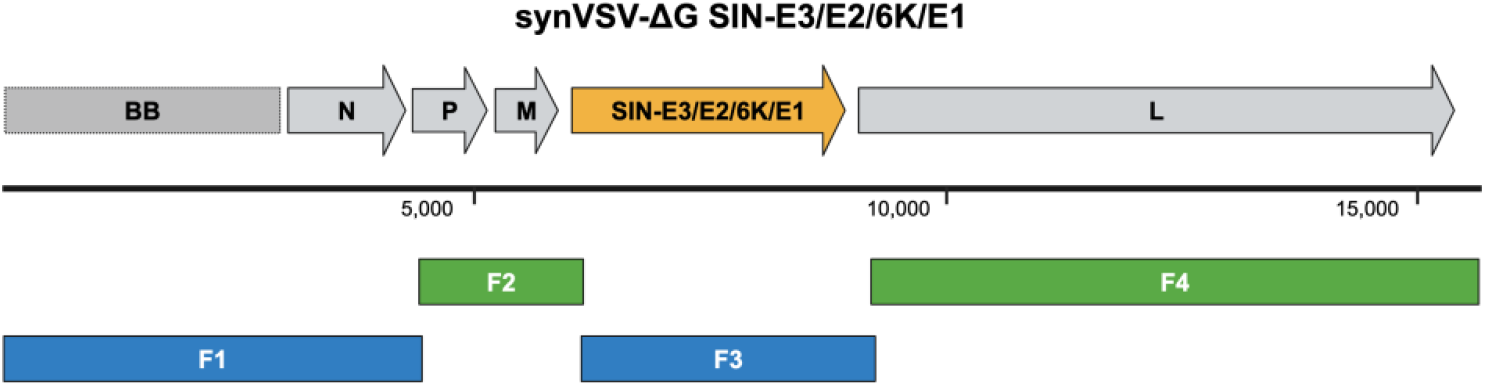
Genome schematic of synVSV-ΔG SIN-E3/E2/6K/E1 is depicted on top. The grey genes represent the normal genes at their standard location. Colored genes represent a change in the genome. Genome size is indicated by the bar below (number is in reference to base pairs). Below in blue, the four fragments used to assemble this construct are shown.

Infectivity assays revealed a distinct difference in the susceptibility of HepaRG and HepG2 cells to synVSV-ΔG SIN-E3/E2/6K/E1 (Fig 6). Notably, minimal infection was observed in HepaRG cells, representing differentiated normal human hepatocyte-like cells, even at a multiplicity of infection (MOI) of 1 after 24 h. In contrast, HepG2 cells, a liver cancer line known for its elevated expression of laminin receptor (LAMR), showed substantial susceptibility to the virus. At an MOI of 0.1, 80% of HepG2 cells were infected within 24 h, while an MOI of 0.01 resulted in 31% infected cells. Interestingly, an MOI of 10 was required to detect infectivity in HepaRG cells, indicating a significantly higher resistance to infection compared to HepG2 cells. These results underscore the preferential tropism of the virus for cancerous liver cells over normal hepatocyte-like cells. To this end, these promising preliminary results warrant further investigation of the oncolytic virus as a potential therapy for liver cancer. Finally, the successful design, construction, and evaluation of synVSV-ΔG SIN-E3/E2/6K/E1 is validation that the synthetic virology platform is suitable for VSV retargeting and evaluating foreign glycoproteins.

**Figure 6.**
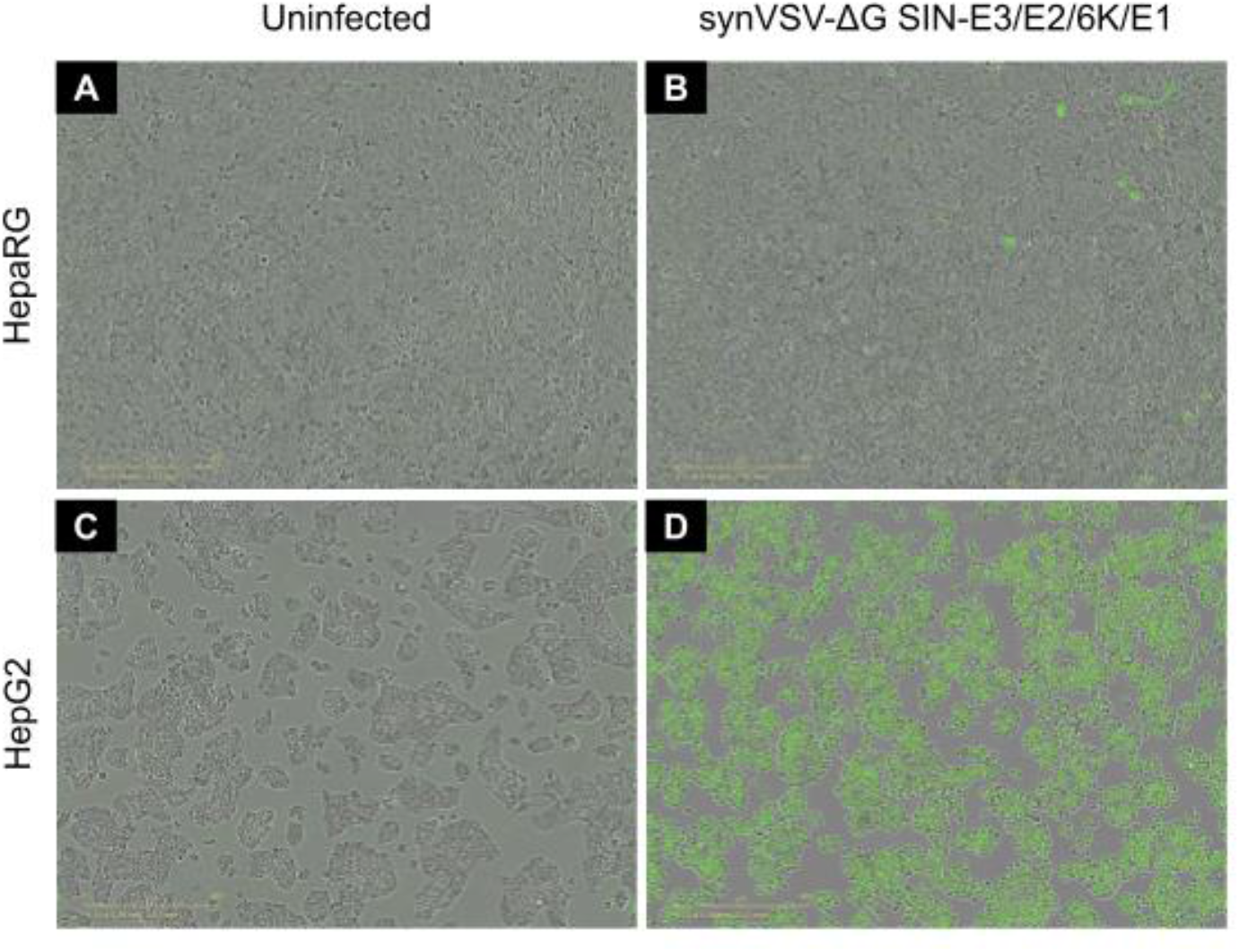
synVSV-ΔG SIN-E3/E2/6K/E1 demonstrated minimal infectivity of HepaRG (differentiated normal, healthy human hepatocyte-like cells) at MOI = 1 after 24 h (**A**,**B**). HepG2 are liver cancer cells that have been reported to have high expression of laminin receptor (LAMR), which is the proposed target receptor for Sindbis virus. In contrast to the normal liver cells, this oncolytic virus exhibited preferential infection of HepG2 cells at the same input and timeline (**C**,**D**).

### 3.4. Evaluate VSV with Gene Rearrangement Using the Synthetic Virology Platform

In this study, we aimed to explore genomic modifications beyond traditional glycoprotein engineering, focusing on how the order and location of VSV genes impact viral fitness and kinetics.

VSV has a negative-sense RNA genome, with gene expression regulated by the viral RNA-dependent RNA polymerase, which initiates transcription at a single promoter near the 3′ end of the RNA and proceeds sequentially. The typical VSV gene order is 3′-N-P-M-G-L-5′. This gene order is highly conserved, with genes encoding proteins required in large amounts, such as the nucleocapsid protein (N), positioned near the promoter, while genes encoding proteins needed in smaller quantities, like the large subunit (L) of the polymerase, are located distally.

Previous studies have shown that rearranging the VSV-N gene attenuates VSV replication and fitness [60] we found few articles addressing other gene rearrangements. However, one study by [61] examined the phenotypic consequences of rearranging the VSV-P, M, and G genes. They found that after 30 h, VSV with the MPG gene rearrangement produced larger plaques than wild-type VSV. Since strain differences, cell type, and other variables were controlled, this greater plaque diameter may indicate a higher viral replication rate, increased cell-to-cell spread, and greater cytopathogenicity. Yang et al. [62], evaluated rearrangement of rabies virus, moving the M gene to the second position (N1M2P3), and found slower replication and attenuated effects. Considering that one of the limitations of oncolytic virus therapies is the reduced efficacy following engineering (e.g., incorporation of promoters, miRNA elements, or foreign glycoproteins), understanding how to enhance these factors could be valuable. To our knowledge, no follow-up studies have been conducted to confirm or refute these findings. Therefore, we decided to validate these observations using our platform.

To test the impact of gene rearrangement, we designed and constructed a VSV with the P and M genes locations changed. The overlap sites at the VSV-N and downstream VSV-G junction/adapter sequences were preserved, and a DNA fragment was synthesized with the natural P-M orientation swapped to M-P. We designated wild-type VSV as VSV-P2M3 (assigning numbers based on gene location in the genome) and named the new construct synVSV-M2P3. Notably, the GT dinucleotide typically found between the P and M genes was maintained. The genome sequences and corresponding fragments for this HiFi assembly are detailed in Supplementary Table S3 and depicted schematically in Figure 7.

**Figure 7.**
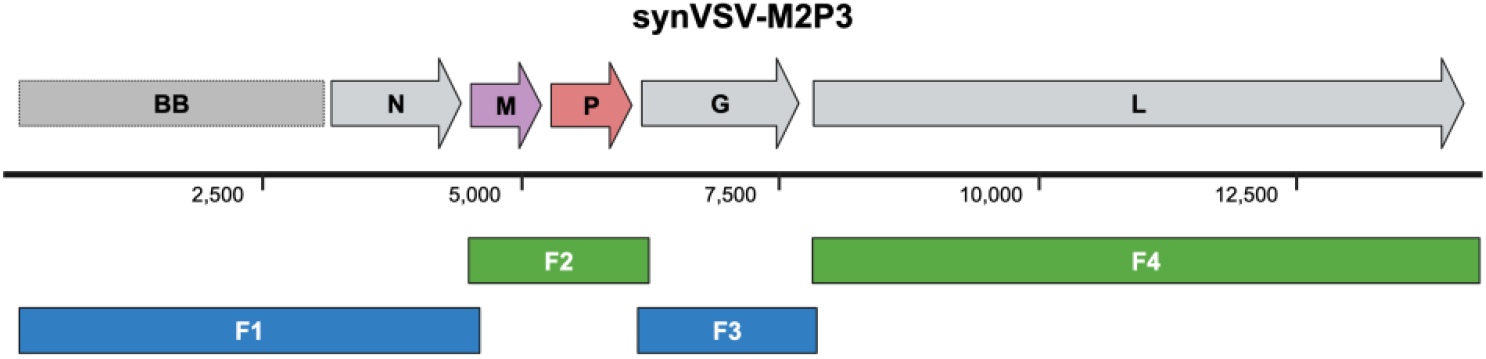
Genome schematic of synVSV-M2P3 is depicted on top. The grey genes represent the normal genes at their standard location. Colored genes represent a change in the genome. Genome size is indicated by the bar below (number is in reference to base pairs). Below in blue, the four fragments used to assemble this construct are shown.

Following genome assembly and sequence validation, synVSV-M2P3 was successfully recovered through reverse genetics, and complete cytopathic effects (CPE) and cell lysis were observed as early as 12–18 h post-infection in passage 1 on 293T cells. Virus recovery was confirmed by RT-qPCR using VSV-M-specific primers. Amplification in 293T cells yielded supernatants with 1.4 × 10^11^ ± 1.1 × 108 genomes/mL and an average functional titer of 3.2 × 10^9^ TCID50/mL, as determined by duplicate TCID_50_ assays.

Subsequent serial infections of both synVSV and synVSV-M2P3 on 293T (production cells) and Hep3B (liver cancer cells) were conducted at various MOIs, with real-time monitoring via the Incucyte system. Both viruses demonstrated rapid onset of CPE and lysis, but as the assay progressed, differences in replication kinetics and potency became apparent, particularly at lower MOIs. After 72 h, immunocytochemistry (ICC) was performed, and phase and green fluorescence (AlexaFluor-488) images (10×) were captured for analysis. Endpoint data showed that on 293T cells, a MOI of 0.0005 (95% CI: 0.0001 to 0.001) resulted in 50% cell confluence inhibition or reduction relative to untreated, while a MOI of 0.003 (95% CI: 0.001 to 0.008) was required to achieve the same effect for synVSV-M2P3. (Figure 8A). On Hep3B cells, synVSV demonstrated 50% reduction with a MOI of 0.002 (95% CI: 1.687 × 10^−5^ to 0.051), and synVSV-M2P3 needed an MOI of 0.033 (95% CI: 0.004 to 6.511) for equivalent effects (Figure 8B).

**Figure 8.**
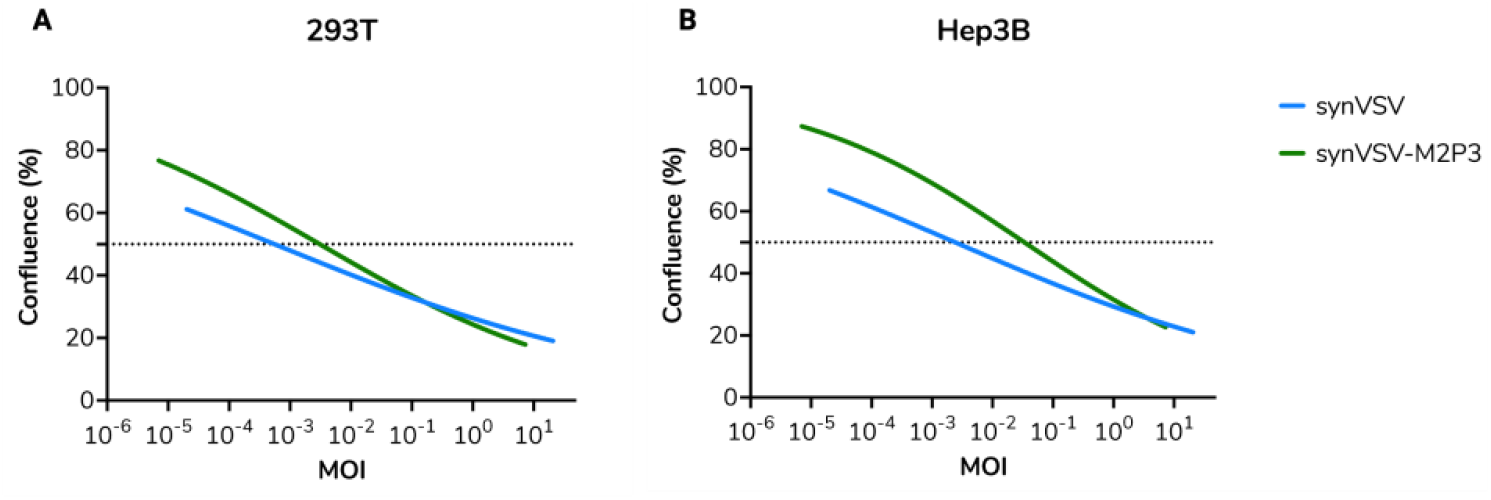
synVSV and synVSV-M2P3 were evaluated for viral kinetics on 293T (**A**) and Hep3B (**B**). Serial ten-fold dilutions starting at MOI = 10 to 10^−5^ were evaluated. For cell health and proliferation evaluation, images were analyzed for cell confluence (%) every 6 h for over 24 h. At the endpoint, confluence was plotted and assessed here. The dashed line represents 50% confluence relative to uninfected wells at endpoint.

These results indicate that synVSV-M2P3 shows comparable lytic activity at high MOIs, but this advantage diminishes at lower viral inputs, where the modified virus demonstrates slower kinetics compared to wild-type VSV. While it is possible that Ball et al. observed larger plaques from VSV-M2P3, this assay and findings align with previous reports of gene rearrangement leading to attenuation. Beyond assessing the impact of gene rearrangement on VSV kinetics, these results further validate the synthetic virology platform described in this study, demonstrating the efficiency of nontraditional modifications. Specifically, the M-P gene rearrangement was achieved within just 5 days (~30 min for design, 3 days for DNA synthesis by Twist Bioscience, and 2 days for assembly, transformation, and validation of the construct), while virus recovery and verification were completed in a total of 9 days, including transfection, amplification, and confirmation by qPCR and nanopore sequencing. This rapid timeline highlights the platform’s potential for accelerating viral engineering.

## 4. Discussion

Here, we demonstrate a de novo approach to the construction of VSV, which represents a technical advance in speed and versatility of generated VSV-based vectors. Modular assembly of DNA viruses [63] and versatile reverse genetics platforms have been reported for positive-strand RNA viruses, especially for SARS-CoV-2[56, 64, 65]. However, the application and benefits of these methods have not been reported for VSV, nor negative-strand RNA viruses for that matter as far as we are currently aware.

Traditional virus engineering techniques, such as recombinant DNA technology and PCR amplification, are often beset by time-consuming processes and susceptibility to errors. Furthermore, traditional molecular biology methods necessitate the availability and use of a DNA template, but de novo synthesis and assembly methods are not bound by this requirement. Thus, synthetic virology methods may be used to (re)construct viruses without available sequence data, extinct pathogens, or entirely new viruses that build upon nature’s design[27]. Taken together, the greatest benefit of building from scratch is the customization of genetic code and complete control over the final construct. To this end, there is an unrealized opportunity to leverage this approach to enable rapid, efficient modifications of VSV genomes.

Other groups have reported protocols to generate recombinant VSV vectors using PCR-based amplification of target genes and then plasmid assembly[23]. A major concern with this approach is that it depends on existing plasmid or available DNA to be able to be amplified before being incorporated into the VSV genome. If research groups are attempting to study outbreak viruses and new strains of concern, the availability of template plasmids is new/limited, which in turn would hamper efforts to study these viruses and their glycoproteins in a safer context. Here, the availability of the digital sequence or shared reference online is the only prerequisite for design and assembly.

In this study, our strategy was to split the VSV genome reference sequence into a set of fragments for DNA synthesis that could be assembled with high-fidelity. The rationale for this approach is that synthetic DNA offers several advantages over more traditional PCR amplification approaches, these include lower error rates, freedom of design or customization (more straightforward mutagenesis, obtain desired sequences with high GC content or repeats that may otherwise be difficult for PCR), ability to obtain longer contiguous sequences or large constructs, purity (e.g., no primer contamination, heterogenous or incomplete amplified fragments).

Within synthetic biology, there is one key difference in our approach compared to other assembly methods is that we only used linear DNA fragments here. Most often, articles that use HiFi Assembly master mix and cloning kit (NEB) have a plasmid backbone that needs to be opened up or linearized (during the reaction) for the assembly. By starting with only linear DNA fragments, the only colonies to pick after transformation are theoretically the correct construct. As bacteria do not uptake linear DNA readily and there are no other circular constructs as potential contaminants, only the desired assembled circular plasmid is available. Again, this is said theoretically, as satellite plasmids are possible or other issues may have occurred (if antibiotic is not added, outdated, etc.). In our study we demonstrated 10/10 colonies to be correct in our screening of the resultant clones.

While there are multiple benefits and advantages to this synthetic virology approach, there are also several potential issues and disadvantages that should be highlighted. First, one drawback of this approach is that the reference sequence must be known, and it must be correct. Public databases and reference sequences are ideally correct, but mutations, sequencing errors and artifacts, assembly and annotation errors, genomic contamination, human error, and other factors are real issues for the scientific community that have serious implications and unintended consequences.

Highlighting just how pervasive of an issue this is, one study found 712 of over 3400 papers analyzed had errors in sequences reported[66]. VectorBuilder, a cloning service provider, reported in a preprint recently that 35% of plasmids (91 of 259 total) they received had sequence variations and differed from the sender/customers’ reference sequence. We experienced this issue firsthand during our first attempts to design and rescue a synthetic VSV.

Designs for synthetic VSV (synVSV) were completed based on the reference file provided (Supplementary Materials, Reference Sequence S2). DNA fragments were synthesized and assembled as described previously, but no virus was recovered after six independent attempted transfections. While troubleshooting, VSV-FL(2+) plasmid was sequenced by Plasmidsaurus, and we learned there were several inconsistencies between the shared reference sequence and actual sequence of the VSV antigenome plasmid. Whole-plasmid sequencing identified 14 nucleotide changes between references. Together, these contributed to one mutation in VSV-N (V14I), two mutations in VSV-P (P77Q, Q110P), two mutations in VSV-M (A133T, G226S), and two mutations in VSV-G (L57I, H96Q), and one mutation in VSV-L (S689T). Based on the new reference sequence, the process was repeated and rescue of synVSV was successful on the first attempt. Of note, this is not a call out against provider Kerafast, as there has been considerable advances in NGS technology since the sequence was deposited. Rather, the aim is to share how commonplace errors are, the need to verify published sequences, and the potential impact these have on synthetic virology and biology.

This de novo synthesis and genome assembly of VSV and variants was a proof-of-concept for the approach. The benefits were realized in these experiments. With the modularized larger fragments in house, the design and synthesis of MP fragment was completed in 3 days. The time from start to virus rescue and sequence confirmation was as little as 4 days. While these benefits were realized, the methods described here can be improved further still. If the VSV genome could be deconstructed into even smaller pieces (5–8 fragments), then new fragments would similarly be smaller, which means higher success rate and shorter timelines from DNA synthesis providers. Optimized overlaps would help facilitate this transition. Ideally, overlaps are 25–30 bp long, GC content of 40–60%, and melting temperature of approximately 50–60 °C. This is a stepwise improvement on the reported methods. More drastic improvements could be achieved by transitioning to using Golden Gate Assembly, as some studies have reported successful assembly of genomes using up to 52 DNA fragments with this approach [67]. To this end, our team has completed initial designs for using this assembly approach, but synthesis is in progress and thus, virus rescue has not been attempted yet. The results of these experiments will be shared in the future when completed. These collective results and potential improvements will lead to the development of novel VSV constructs and therapies with even greater promise to improve human health[68, 69].

## 5. Conclusions

In summary, a synthetic vesicular stomatitis virus (synVSV) was designed, assembled, and rescued. There was no significant difference between synVSV and its natural counterpart. A synthetic virology platform enabled rapid construction and reverse genetics to study this virus. The platform was further validated by producing VSV variants, including i) VSV with VSV-G sequence deleted, and replaced with Sindbis virus envelope glycoprotein, and ii) VSV with the phosphoprotein (VSV-P) and matrix (VSV-M) genes rearranged with VSV-M2P3 order. Taking everything into consideration, synthetic virology enables the rapid engineering of VSV for basic science and molecular virology as well as the development of novel vaccines and therapies with the potential to improve human health.

## 6. Dual Use Research Concerns (DURC)

Beyond the therapeutic benefits and technical advantages of our approach, there are several dual-use research concerns. Chief among these, the development of artificial viruses and/or use of synthetic genomics methods to (re)create select agents. To this end, the capacity to use synthetic DNA to build a viral genome has existed for some time. Commercial DNA synthesis companies adhere to screening guidance and documents that examine orders for legitimacy or nefarious intent. Ultimately, we are aware of and understand the dual-use concerns surrounding this technology, but we believe these are far outweighed by the potential benefits.

## Supporting information

Supplemental Sequence 1

Supplemental Sequence 2

## Acknowledgements

We are grateful to Dr. Michael A. Whitt for all his support and advice on vesicular stomatitis virus, and feedback on this article.

## Supplementary Materials

Reference Sequence S1: VSV wildtype reference sequence. Reference Sequence S2: VSV wildtype reference sequence supplied by Kerafast. Table S1: DNA fragment sequences for genome assembly of synVSV. Table S2: DNA fragment sequences for genome assembly of synVSV-G.Table S2: DNA fragment sequences for genome assembly of SIN-E3/E2/6K/E1. Table S3: DNA fragment sequences for genome assembly of synVSV-M2P3.

## Author Contributions

Conceptualization, C.M.M., R.B. and P.W.; methodology, C.M.M., R.B., P.W., B.F.S.; software, P.W.; validation, R.B., B.H., M.F. and T.F.; formal analysis, C.M.M., R.B., P.W., B.H., M.F., T.F., B.F.S.; resources, P.W. and B.F.S.; data curation, C.M.M., R.B., B.H., M.F. and T.F.; writing—original draft preparation, C.M.M. and R.B.; writing—review and editing, C.M.M., R.B., P.W., B.F.S.; visualization, C.M.M. and R.B.; supervision, C.M.M., R.B. and P.W. All authors have read and agreed to the published version of the manuscript.

## Institutional Review Board Statement

This study does not involve human subjects. Therefore, approval from an Institutional Review Board (IRB) is not applicable or required.

## Data Availability Statement

Supporting data is available in the Supplementary materials.

