## Supplemental Sequence 1 for "Leveraging Synthetic Virology for the Rapid Engineering of Vesicular Stomatitis Virus (VSV)"

TAATACGACTCACTATAGGACGAAGACAAACAAACCATTATTATCATTAAAAGGCTCAGGAGAAACTTTAACAGTAATCAAAATGTCTGTTACAGTCAAGAGAATCATTGACAACACAGTCGTAGTTCCAAAACTTCCTGCAAATGAGGATCCAGTGGAATACCCGGCAGATTACTTCAGAAAATCAAAGGAGATTCCTCTTTACATCAATACTACAAAAAGTTTGTCAGATCTAAGAGGATATGTCTACCAAGGCCTCAAATCCGGAAATGTATCAATCATACATGTCAACAGCTACTTGTATGGAGCATTAAAGGACATCCGGGGTAAGTTGGATAAAGATTGGTCAAGTTTCGGAATAAACATCGGGAAAGCAGGGGATACAATCGGAATATTTGACCTTGTATCCTTGAAAGCCCTGGACGGCGTACTTCCAGATGGAGTATCGGATGCTTCCAGAACCAGCGCAGATGACAAATGGTTGCCTTTGTATCTACTTGGCTTATACAGAGTGGGCAGAACACAAATGCCTGAATACAGAAAAAAGCTCATGGATGGGCTGACAAATCAATGCAAAATGATCAATGAACAGTTTGAACCTCTTGTGCCAGAAGGTCGTGACATTTTTGATGTGTGGGGAAATGACAGTAATTACACAAAAATTGTCGCTGCAGTGGACATGTTCTTCCACATGTTCAAAAAACATGAATGTGCCTCGTTCAGATACGGAACTATTGTTTCCAGATTCAAAGATTGTGCTGCATTGGCAACATTTGGACACCTCTGCAAAATAACCGGAATGTCTACAGAAGATGTAACGACCTGGATCTTGAACCGAGAAGTTGCAGATGAAATGGTCCAAATGATGCTTCCAGGCCAAGAAATTGACAAGGCCGATTCATACATGCCTTATTTGATCGACTTTGGATTGTCTTCTAAGTCTCCATATTCTTCCGTCAAAAACCCTGCCTTCCACTTCTGGGGGCAATTGACAGCTCTTCTGCTCAGATCCACCAGAGCAAGGAATGCCCGACAGCCTGATGACATTGAGTATACATCTCTTACTACAGCAGGTTTGTTGTACGCTTATGCAGTAGGATCCTCTGCCGACTTGGCACAACAGTTTTGTGTTGGAGATAACAAATACACTCCAGATGATAGTACCGGAGGATTGACGACTAATGCACCGCCACAAGGCAGAGATGTGGTCGAATGGCTCGGATGGTTTGAAGATCAAAACAGAAAACCGACTCCTGATATGATGCAGTATGCGAAAAGAGCAGTCATGTCACTGCAAGGCCTAAGAGAGAAGACAATTGGCAAGTATGCTAAGTCAGAATTTGACAAATGACCCTATAATTCTCAGATCACCTATTATATATTATGCTACATATGAAAAAAACTAACAGATATCATGGATAATCTCACAAAAGTTCGTGAGTATCTCAAGTCCTATTCTCGTCTGGATCAGGCGGTAGGAGAGATAGATGAGATCGAAGCACAACGAGCTGAAAAGTCCAATTATGAGTTGTTCCAAGAGGATGGAGTGGAAGAGCATACTAAGCCCTCTTATTTTCAGGCAGCAGATGATTCTGACACAGAATCTGAACCAGAAATTGAAGACAATCAAGGCTTGTATGCACCAGATCCAGAAGCTGAGCAAGTTGAAGGCTTTATACAGGGGCCTTTAGATGACTATGCAGATGAGGAAGTGGATGTTGTATTTACTTCGGACTGGAAACAGCCTGAGCTTGAATCTGACGAGCATGGAAAGACCTTACGGTTGACATCGCCAGAGGGTTTAAGTGGAGAGCAGAAATCCCAGTGGCTTTCGACGATTAAAGCAGTCGTGCAAAGTGCCAAATACTGGAATCTGGCAGAGTGCACATTTGAAGCATCGGGAGAAGGGGTCATTATGAAGGAGCGCCAGATAACTCCGGATGTATATAAGGTCACTCCAGTGATGAACACACATCCGTCCCAATCAGAAGCAGTATCAGATGTTTGGTCTCTCTCAAAGACATCCATGACTTTCCAACCCAAGAAAGCAAGTCTTCAGCCTCTCACCATATCCTTGGATGAATTGTTCTCATCTAGAGGAGAGTTCATCTCTGTCGGAGGTGACGGACGAATGTCTCATAAAGAGGCCATCCTGCTCGGCCTGAGATACAAAAAGTTGTACAATCAGGCGAGAGTCAAATATTCTCTGTAGACTATGAAAAAAAGTAACAGATATCACGATCTAAGTGTTATCCCAATCCATTCATCATGAGTTCCTTAAAGAAGATTCTCGGTCTGAAGGGGAAAGGTAAGAAATCTAAGAAATTAGGGATCGCACCACCCCCTTATGAAGAGGACACTAGCATGGAGTATGCTCCGAGCGCTCCAATTGACAAATCCTATTTTGGAGTTGACGAGATGGACACCTATGATCCGAATCAATTAAGATATGAGAAATTCTTCTTTACAGTGAAAATGACGGTTAGATCTAATCGTCCGTTCAGAACATACTCAGATGTGGCAGCCGCTGTATCCCATTGGGATCACATGTACATCGGAATGGCAGGGAAACGTCCCTTCTACAAAATCTTGGCTTTTTTGGGTTCTTCTAATCTAAAGGCCACTCCAGCGGTATTGGCAGATCAAGGTCAACCAGAGTATCACGCTCACTGCGAAGGCAGGGCTTATTTGCCACATAGGATGGGGAAGACCCCTCCCATGCTCAATGTACCAGAGCACTTCAGAAGACCATTCAATATAGGTCTTTACAAGGGAACGATTGAGCTCACAATGACCATCTACGATGATGAGTCACTGGAAGCAGCTCCTATGATCTGGGATCATTTCAATTCTTCCAAATTTTCTGATTTCAGAGAGAAGGCCTTAATGTTTGGCCTGATTGTCGAGAAAAAGGCATCTGGAGCGTGGGTCCTGGACTCTATCGGCCACTTCAAATGAGCTAGTCTAACTTCTAGCTTCTGAACAATCCCCGGTTTACTCAGTCTCCCCTAATTCCAGCCTCTCGAACAACTAATATCCTGTCTTTTCTATCCCTATGAAAAAAACTAACAGAGATCGATCTGTTTACGCGTCACTATGAAGTGCCTTTTGTACTTAGCCTTTTTATTCATTGGGGTGAATTGCAAGTTCACCATAGTTTTTCCACACAACCAAAAAGGAAACTGGAAAAATGTTCCTTCTAATTACCATTATTGCCCGTCAAGCTCAGATTTAAATTGGCATAATGACTTAATAGGCACAGCCTTACAAGTCAAAATGCCCAAGAGTCACAAGGCTATTCAAGCAGACGGTTGGATGTGTCATGCTTCCAAATGGGTCACTACTTGTGATTTCCGCTGGTATGGACCGAAGTATATAACACATTCCATCCGATCCTTCACTCCATCTGTAGAACAATGCAAGGAAAGCATTGAACAAACGAAACAAGGAACTTGGCTGAATCCAGGCTTCCCTCCTCAAAGTTGTGGATATGCAACTGTGACGGATGCCGAAGCAGTGATTGTCCAGGTGACTCCTCACCATGTGCTGGTTGATGAATACACAGGAGAATGGGTTGATTCACAGTTCATCAACGGAAAATGCAGCAATTACATATGCCCCACTGTCCATAACTCTACAACCTGGCATTCTGACTATAAGGTCAAAGGGCTATGTGATTCTAACCTCATTTCCATGGACATCACCTTCTTCTCAGAGGACGGAGAGCTATCATCCCTGGGAAAGGAGGGCACAGGGTTCAGAAGTAACTACTTTGCTTATGAAACTGGAGGCAAGGCCTGCAAAATGCAATACTGCAAGCATTGGGGAGTCAGACTCCCATCAGGTGTCTGGTTCGAGATGGCTGATAAGGATCTCTTTGCTGCAGCCAGATTCCCTGAATGCCCAGAAGGGTCAAGTATCTCTGCTCCATCTCAGACCTCAGTGGATGTAAGTCTAATTCAGGACGTTGAGAGGATCTTGGATTATTCCCTCTGCCAAGAAACCTGGAGCAAAATCAGAGCGGGTCTTCCAATCTCTCCAGTGGATCTCAGCTATCTTGCTCCTAAAAACCCAGGAACCGGTCCTGCTTTCACCATAATCAATGGTACCCTAAAATACTTTGAGACCAGATACATCAGAGTCGATATTGCTGCTCCAATCCTCTCAAGAATGGTCGGAATGATCAGTGGAACTACCACAGAAAGGGAACTGTGGGATGACTGGGCACCATATGAAGACGTGGAAATTGGACCCAATGGAGTTCTGAGGACCAGTTCAGGATATAAGTTTCCTTTATACATGATTGGACATGGTATGTTGGACTCCGATCTTCATCTTAGCTCAAAGGCTCAGGTGTTCGAACATCCTCACATTCAAGACGCTGCTTCGCAACTTCCTGATGATGAGAGTTTATTTTTTGGTGATACTGGGCTATCCAAAAATCCAATCGAGCTTGTAGAAGGTTGGTTCAGTAGTTGGAAAAGCTCTATTGCCTCTTTTTTCTTTATCATAGGGTTAATCATTGGACTATTCTTGGTTCTCCGAGTTGGTATCCATCTTTGCATTAAATTAAAGCACACCAAGAAAAGACAGATTTATACAGACATAGAGATGAACCGACTTGGAAAGTAACTCAAATCCTGCTAGCCAGATTCTTCATGTTTGGACCAAATCAACTTGTGATACCATGCTCAAAGAGGCCTCAATTATATTTGAGTTTTTAATTTTTATGAAAAAAACTAACAGCAATCATGGAAGTCCACGATTTTGAGACCGACGAGTTCAATGATTTCAATGAAGATGACTATGCCACAAGAGAATTCCTGAATCCCGATGAGCGCATGACGTACTTGAATCATGCTGATTACAACCTGAATTCTCCTCTAATTAGTGATGATATTGACAATTTAATCAGGAAATTCAATTCTCTTCCAATTCCCTCGATGTGGGATAGTAAGAACTGGGATGGAGTTCTTGAGATGTTAACATCATGTCAAGCCAATCCCATCTCAACATCTCAGATGCATAAATGGATGGGAAGTTGGTTAATGTCTGATAATCATGATGCCAGTCAAGGGTATAGTTTTTTACATGAAGTGGACAAAGAGGCAGAAATAACATTTGACGTGGTGGAGACCTTCATCCGCGGCTGGGGCAACAAACCAATTGAATACATCAAAAAGGAAAGATGGACTGACTCATTCAAAATTCTCGCTTATTTGTGTCAAAAGTTTTTGGACTTACACAAGTTGACATTAATCTTAAATGCTGTCTCTGAGGTGGAATTGCTCAACTTGGCGAGGACTTTCAAAGGCAAAGTCAGAAGAAGTTCTCATGGAACGAACATATGCAGGATTAGGGTTCCCAGCTTGGGTCCTACTTTTATTTCAGAAGGATGGGCTTACTTCAAGAAACTTGATATTCTAATGGACCGAAACTTTCTGTTAATGGTCAAAGATGTGATTATAGGGAGGATGCAAACGGTGCTATCCATGGTATGTAGAATAGACAACCTGTTCTCAGAGCAAGACATCTTCTCCCTTCTAAATATCTACAGAATTGGAGATAAAATTGTGGAGAGGCAGGGAAATTTTTCTTATGACTTGATTAAAATGGTGGAACCGATATGCAACTTGAAGCTGATGAAATTAGCAAGAGAATCAAGGCCTTTAGTCCCACAATTCCCTCATTTTGAAAATCATATCAAGACTTCTGTTGATGAAGGGGCAAAAATTGACCGAGGTATAAGATTCCTCCATGATCAGATAATGAGTGTGAAAACAGTGGATCTCACACTGGTGATTTATGGATCGTTCAGACATTGGGGTCATCCTTTTATAGATTATTACACTGGACTAGAAAAATTACATTCCCAAGTAACCATGAAGAAAGATATTGATGTGTCATATGCAAAAGCACTTGCAAGTGATTTAGCTCGGATTGTTCTATTTCAACAGTTCAATGATCATAAAAAGTGGTTCGTGAATGGAGACTTGCTCCCTCATGATCATCCCTTTAAAAGTCATGTTAAAGAAAATACATGGCCCACAGCTGCTCAAGTTCAAGATTTTGGAGATAAATGGCATGAACTTCCGCTGATTAAATGTTTTGAAATACCCGACTTACTAGACCCATCGATAATATACTCTGACAAAAGTCATTCAATGAATAGGTCAGAGGTGTTGAAACATGTCCGAATGAATCCGAACACTCCTATCCCTAGTAAAAAGGTGTTGCAGACTATGTTGGACACAAAGGCTACCAATTGGAAAGAATTTCTTAAAGAGATTGATGAGAAGGGCTTAGATGATGATGATCTAATTATTGGTCTTAAAGGAAAGGAGAGGGAACTGAAGTTGGCAGGTAGATTTTTCTCCCTAATGTCTTGGAAATTGCGAGAATACTTTGTAATTACCGAATATTTGATAAAGACTCATTTCGTCCCTATGTTTAAAGGCCTGACAATGGCGGACGATCTAACTGCAGTCATTAAAAAGATGTTAGATTCCTCATCCGGCCAAGGATTGAAGTCATATGAGGCAATTTGCATAGCCAATCACATTGATTACGAAAAATGGAATAACCACCAAAGGAAGTTATCAAACGGCCCAGTGTTCCGAGTTATGGGCCAGTTCTTAGGTTATCCATCCTTAATCGAGAGAACTCATGAATTTTTTGAGAAAAGTCTTATATACTACAATGGAAGACCAGACTTGATGCGTGTTCACAACAACACACTGATCAATTCAACCTCCCAACGAGTTTGTTGGCAAGGACAAGAGGGTGGACTGGAAGGTCTACGGCAAAAAGGATGGAGTATCCTCAATCTACTGGTTATTCAAAGAGAGGCTAAAATCAGAAACACTGCTGTCAAAGTCTTGGCACAAGGTGATAATCAAGTTATTTGCACACAGTATAAAACGAAGAAATCGAGAAACGTTGTAGAATTACAGGGTGCTCTCAATCAAATGGTTTCTAATAATGAGAAAATTATGACTGCAATCAAAATAGGGACAGGGAAGTTAGGACTTTTGATAAATGACGATGAGACTATGCAATCTGCAGATTACTTGAATTATGGAAAAATACCGATTTTCCGTGGAGTGATTAGAGGGTTAGAGACCAAGAGATGGTCACGAGTGACTTGTGTCACCAATGACCAAATACCCACTTGTGCTAATATAATGAGCTCAGTTTCCACAAATGCTCTCACCGTAGCTCATTTTGCTGAGAACCCAATCAATGCCATGATACAGTACAATTATTTTGGGACATTTGCTAGACTCTTGTTGATGATGCATGATCCTGCTCTTCGTCAATCATTGTATGAAGTTCAAGATAAGATACCGGGCTTGCACAGTTCTACTTTCAAATACGCCATGTTGTATTTGGACCCTTCCATTGGAGGAGTGTCGGGCATGTCTTTGTCCAGGTTTTTGATTAGAGCCTTCCCAGATCCCGTAACAGAAAGTCTCTCATTCTGGAGATTCATCCATGTACATGCTCGAAGTGAGCATCTGAAGGAGATGAGTGCAGTATTTGGAAACCCCGAGATAGCCAAGTTTCGAATAACTCACATAGACAAGCTAGTAGAAGATCCAACCTCTCTGAACATCGCTATGGGAATGAGTCCAGCGAACTTGTTAAAGACTGAGGTTAAAAAATGCTTAATCGAATCAAGACAAACCATCAGGAACCAGGTGATTAAGGATGCAACCATATATTTGTATCATGAAGAGGATCGGCTCAGAAGTTTCTTATGGTCAATAAATCCTCTGTTCCCTAGATTTTTAAGTGAATTCAAATCAGGCACTTTTTTGGGAGTCGCAGACGGGCTCATCAGTCTATTTCAAAATTCTCGTACTATTCGGAACTCCTTTAAGAAAAAGTATCATAGGGAATTGGATGATTTGATTGTGAGGAGTGAGGTATCCTCTTTGACACATTTAGGGAAACTTCATTTGAGAAGGGGATCATGTAAAATGTGGACATGTTCAGCTACTCATGCTGACACATTAAGATACAAATCCTGGGGCCGTACAGTTATTGGGACAACTGTACCCCATCCATTAGAAATGTTGGGTCCACAACATCGAAAAGAGACTCCTTGTGCACCATGTAACACATCAGGGTTCAATTATGTTTCTGTGCATTGTCCAGACGGGATCCATGACGTCTTTAGTTCACGGGGACCATTGCCTGCTTATCTAGGGTCTAAAACATCTGAATCTACATCTATTTTGCAGCCTTGGGAAAGGGAAAGCAAAGTCCCACTGATTAAAAGAGCTACACGTCTTAGAGATGCTATCTCTTGGTTTGTTGAACCCGACTCTAAACTAGCAATGACTATACTTTCTAACATCCACTCTTTAACAGGCGAAGAATGGACCAAAAGGCAGCATGGGTTCAAAAGAACAGGGTCTGCCCTTCATAGGTTTTCGACATCTCGGATGAGCCATGGTGGGTTCGCATCTCAGAGCACTGCAGCATTGACCAGGTTGATGGCAACTACAGACACCATGAGGGATCTGGGAGATCAGAATTTCGACTTTTTATTCCAAGCAACGTTGCTCTATGCTCAAATTACCACCACTGTTGCAAGAGACGGATGGATCACCAGTTGTACAGATCATTATCATATTGCCTGTAAGTCCTGTTTGAGACCCATAGAAGAGATCACCCTGGACTCAAGTATGGACTACACGCCCCCAGATGTATCCCATGTGCTGAAGACATGGAGGAATGGGGAAGGTTCGTGGGGACAAGAGATAAAACAGATCTATCCTTTAGAAGGGAATTGGAAGAATTTAGCACCTGCTGAGCAATCCTATCAAGTCGGCAGATGTATAGGTTTTCTATATGGAGACTTGGCGTATAGAAAATCTACTCATGCCGAGGACAGTTCTCTATTTCCTCTATCTATACAAGGTCGTATTAGAGGTCGAGGTTTCTTAAAAGGGTTGCTAGACGGATTAATGAGAGCAAGTTGCTGCCAAGTAATACACCGGAGAAGTCTGGCTCATTTGAAGAGGCCGGCCAACGCAGTGTACGGAGGTTTGATTTACTTGATTGATAAATTGAGTGTATCACCTCCATTCCTTTCTCTTACTAGATCAGGACCTATTAGAGACGAATTAGAAACGATTCCCCACAAGATCCCAACCTCCTATCCGACAAGCAACCGTGATATGGGGGTGATTGTCAGAAATTACTTCAAATACCAATGCCGTCTAATTGAAAAGGGAAAATACAGATCACATTATTCACAATTATGGTTATTCTCAGATGTCTTATCCATAGACTTCATTGGACCATTCTCTATTTCCACCACCCTCTTGCAAATCCTATACAAGCCATTTTTATCTGGGAAAGATAAGAATGAGTTGAGAGAGCTGGCAAATCTTTCTTCATTGCTAAGATCAGGAGAGGGGTGGGAAGACATACATGTGAAATTCTTCACCAAGGACATATTATTGTGTCCAGAGGAAATCAGACATGCTTGCAAGTTCGGGATTGCTAAGGATAATAATAAAGACATGAGCTATCCCCCTTGGGGAAGGGAATCCAGAGGGACAATTACAACAATCCCTGTTTATTATACGACCACCCCTTACCCAAAGATGCTAGAGATGCCTCCAAGAATCCAAAATCCCCTGCTGTCCGGAATCAGGTTGGGCCAATTACCAACTGGCGCTCATTATAAAATTCGGAGTATATTACATGGAATGGGAATCCATTACAGGGACTTCTTGAGTTGTGGAGACGGCTCCGGAGGGATGACTGCTGCATTACTACGAGAAAATGTGCATAGCAGAGGAATATTCAATAGTCTGTTAGAATTATCAGGGTCAGTCATGCGAGGCGCCTCTCCTGAGCCCCCCAGTGCCCTAGAAACTTTAGGAGGAGATAAATCGAGATGTGTAAATGGTGAAACATGTTGGGAATATCCATCTGACTTATGTGACCCAAGGACTTGGGACTATTTCCTCCGACTCAAAGCAGGCTTGGGGCTTCAAATTGATTTAATTGTAATGGATATGGAAGTTCGGGATTCTTCTACTAGCCTGAAAATTGAGACGAATGTTAGAAATTATGTGCACCGGATTTTGGATGAGCAAGGAGTTTTAATCTACAAGACTTATGGAACATATATTTGTGAGAGCGAAAAGAATGCAGTAACAATCCTTGGTCCCATGTTCAAGACGGTCGACTTAGTTCAAACAGAATTTAGTAGTTCTCAAACGTCTGAAGTATATATGGTATGTAAAGGTTTGAAGAAATTAATCGATGAACCCAATCCCGATTGGTCTTCCATCAATGAATCCTGGAAAAACCTGTACGCATTCCAGTCATCAGAACAGGAATTTGCCAGAGCAAAGAAGGTTAGTACATACTTTACCTTGACAGGTATTCCCTCCCAATTCATTCCTGATCCTTTTGTAAACATTGAGACTATGCTACAAATATTCGGAGTACCCACGGGTGTGTCTCATGCGGCTGCCTTAAAATCATCTGATAGACCTGCAGATTTATTGACCATTAGCCTTTTTTATATGGCGATTATATCGTATTATAACATCAATCATATCAGAGTAGGACCGATACCTCCGAACCCCCCATCAGATGGAATTGCACAAAATGTGGGGATCGCTATAACTGGTATAAGCTTTTGGCTGAGTTTGATGGAGAAAGACATTCCACTATATCAACAGTGTTTAGCAGTTATCCAGCAATCATTCCCGATTAGGTGGGAGGCTGTTTCAGTAAAAGGAGGATACAAGCAGAAGTGGAGTACTAGAGGTGATGGGCTCCCAAAAGATACCCGAATTTCAGACTCCTTGGCCCCAATCGGGAACTGGATCAGATCTCTGGAATTGGTCCGAAACCAAGTTCGTCTAAATCCATTCAATGAGATCTTGTTCAATCAGCTATGTCGTACAGTGGATAATCATTTGAAATGGTCAAATTTGCGAAGAAACACAGGAATGATTGAATGGATCAATAGACGAATTTCAAAAGAAGACCGGTCTATACTGATGTTGAAGAGTGACCTACACGAGGAAAACTCTTGGAGAGATTAAAAAATCATGAGGAGACTCCAAACTTTAAGTATGAAAAAAACTTTGATCCTTAAGACCCTCTTGTGGTTTTTATTTTTTATCTGGTTTTGTGGTCTTCGTGGGTCGGCATGGCATCTCCACCTCCTCGCGGTCCGACCTGGGCATCCGAAGGAGGACGTCGTCCACTCGGATGGCTAAGGGAGGGGCCCCCGCGGGGCTGCTAACAAAGCCCGAAAGGAAGCTGAGTTGGCTGCTGCCACCGCTGAGCAATAACTAGCATAACCCCTTGGGGCCTCTAAACGGGTCTTGAGGGGTTTTTTGCTGAAAGGAGGAACTATATCCGGATCGAGACCTCGATACTAGTGCGGTGGAGCTCCAGCTTTTGTTCCCTTTAGTGAGGGTTAATTTCGAGCTTGGCGTAATCATGGTCATAGCTGTTTCCTGTGTGAAATTGTTATCCGCTCACAATTCCACACAACATACGAGCCGGAAGCATAAAGTGTAAAGCCTGGGGTGCCTAATGAGTGAGCTAACTCACATTAATTGCGTTGCGCTCACTGCCCGCTTTCCAGTCGGGAAACCTGTCGTGCCAGCTGCATTAATGAATCGGCCAACGCGCGGGGAGAGGCGGTTTGCGTATTGGGCGCTCTTCCGCTTCCTCGCTCACTGACTCGCTGCGCTCGGTCGTTCGGCTGCGGCGAGCGGTATCAGCTCACTCAAAGGCGGTAATACGGTTATCCACAGAATCAGGGGATAACGCAGGAAAGAACATGTGAGCAAAAGGCCAGCAAAAGGCCAGGAACCGTAAAAAGGCCGCGTTGCTGGCGTTTTTCCATAGGCTCCGCCCCCCTGACGAGCATCACAAAAATCGACGCTCAAGTCAGAGGTGGCGAAACCCGACAGGACTATAAAGATACCAGGCGTTTCCCCCTGGAAGCTCCCTCGTGCGCTCTCCTGTTCCGACCCTGCCGCTTACCGGATACCTGTCCGCCTTTCTCCCTTCGGGAAGCGTGGCGCTTTCTCATAGCTCACGCTGTAGGTATCTCAGTTCGGTGTAGGTCGTTCGCTCCAAGCTGGGCTGTGTGCACGAACCCCCCGTTCAGCCCGACCGCTGCGCCTTATCCGGTAACTATCGTCTTGAGTCCAACCCGGTAAGACACGACTTATCGCCACTGGCAGCAGCCACTGGTAACAGGATTAGCAGAGCGAGGTATGTAGGCGGTGCTACAGAGTTCTTGAAGTGGTGGCCTAACTACGGCTACACTAGAAGAACAGTATTTGGTATCTGCGCTCTGCTGAAGCCAGTTACCTTCGGAAAAAGAGTTGGTAGCTCTTGATCCGGCAAACAAACCACCGCTGGTAGCGGTGGTTTTTTTGTTTGCAAGCAGCAGATTACGCGCAGAAAAAAAGGATCTCAAGAAGATCCTTTGATCTTTTCTACGGGGTCTGACGCTCAGTGGAACGAAAACTCACGTTAAGGGATTTTGGTCATGAGATTATCAAAAAGGATCTTCACCTAGATCCTTTTAAATTAAAAATGAAGTTTTAAATCAATCTAAAGTATATATGAGTAAACTTGGTCTGACAGTTACCAATGCTTAATCAGTGAGGCACCTATCTCAGCGATCTGTCTATTTCGTTCATCCATAGTTGCCTGACTCCCCGTCGTGTAGATAACTACGATACGGGAGGGCTTACCATCTGGCCCCAGTGCTGCAATGATACCGCGAGACCCACGCTCACCGGCTCCAGATTTATCAGCAATAAACCAGCCAGCCGGAAGGGCCGAGCGCAGAAGTGGTCCTGCAACTTTATCCGCCTCCATCCAGTCTATTAATTGTTGCCGGGAAGCTAGAGTAAGTAGTTCGCCAGTTAATAGTTTGCGCAACGTTGTTGCCATTGCTACAGGCATCGTGGTGTCACGCTCGTCGTTTGGTATGGCTTCATTCAGCTCCGGTTCCCAACGATCAAGGCGAGTTACATGATCCCCCATGTTGTGCAAAAAAGCGGTTAGCTCCTTCGGTCCTCCGATCGTTGTCAGAAGTAAGTTGGCCGCAGTGTTATCACTCATGGTTATGGCAGCACTGCATAATTCTCTTACTGTCATGCCATCCGTAAGATGCTTTTCTGTGACTGGTGAGTACTCAACCAAGTCATTCTGAGAATAGTGTATGCGGCGACCGAGTTGCTCTTGCCCGGCGTCAATACGGGATAATACCGCGCCACATAGCAGAACTTTAAAAGTGCTCATCATTGGAAAACGTTCTTCGGGGCGAAAACTCTCAAGGATCTTACCGCTGTTGAGATCCAGTTCGATGTAACCCACTCGTGCACCCAACTGATCTTCAGCATCTTTTACTTTCACCAGCGTTTCTGGGTGAGCAAAAACAGGAAGGCAAAATGCCGCAAAAAAGGGAATAAGGGCGACACGGAAATGTTGAATACTCATACTCTTCCTTTTTCAATATTATTGAAGCATTTATCAGGGTTATTGTCTCATGAGCGGATACATATTTGAATGTATTTAGAAAAATAAACAAATAGGGGTTCCGCGCACATTTCCCCGAAAAGTGCCACCTAAATTGTAAGCGTTAATATTTTGTTAAAATTCGCGTTAAATTTTTGTTAAATCAGCTCATTTTTTAACCAATAGGCCGAAATCGGCAAAATCCCTTATAAATCAAAAGAATAGACCGAGATAGGGTTGAGTGTTGTTCCAGTTTGGAACAAGAGTCCACTATTAAAGAACGTGGACTCCAACGTCAAAGGGCGAAAAACCGTCTATCAGGGCGATGGCCCACTACGTGAACCATCACCCTAATCAAGTTTTTTGGGGTCGAGGTGCCGTAAAGCACTAAATCGGAACCCTAAAGGGAGCCCCCGATTTAGAGCTTGACGGGGAAAGCCGGCGAACGTGGCGAGAAAGGAAGGGAAGAAAGCGAAAGGAGCGGGCGCTAGGGCGCTGGCAAGTGTAGCGGTCACGCTGCGCGTAACCACCACACCCGCCGCGCTTAATGCGCCGCTACAGGGCGCGTCCCATTCGCCATTCAGGCTGCGCAACTGTTGGGAAGGGCGATCGGTGCGGGCCTCTTCGCTATTACGCCAGCTGGCGAAAGGGGGATGTGCTGCAAGGCGATTAAGTTGGGTAACGCCAGGGTTTTCCCAGTCACGACGTTGTAAAACGACGGCCAGTGAATTG
