## Supplemental Table 1 for "Leveraging Synthetic Virology for the Rapid Engineering of Vesicular Stomatitis Virus (VSV)"

| **Fragment #** | **Length (bp)** | **Sequence** |
| --- | --- | --- |
| F1 | 4,450 | GGGTCGGCATGGCATCTCCACCTCCTCGCGGTCCGACCTGGGCATCCGAAGGAGGACGTCGTCCACTCGGATGGCTAAGGGAGGGGCCCCCGCGGGGCTGCTAACAAAGCCCGAAAGGAAGCTGAGTTGGCTGCTGCCACCGCTGAGCAATAACTAGCATAACCCCTTGGGGCCTCTAAACGGGTCTTGAGGGGTTTTTTGCTGAAAGGAGGAACTATATCCGGATCGAGACCTCGATACTAGTGCGGTGGAGCTCCAGCTTTTGTTCCCTTTAGTGAGGGTTAATTTCGAGCTTGGCGTAATCATGGTCATAGCTGTTTCCTGTGTGAAATTGTTATCCGCTCACAATTCCACACAACATACGAGCCGGAAGCATAAAGTGTAAAGCCTGGGGTGCCTAATGAGTGAGCTAACTCACATTAATTGCGTTGCGCTCACTGCCCGCTTTCCAGTCGGGAAACCTGTCGTGCCAGCTGCATTAATGAATCGGCCAACGCGCGGGGAGAGGCGGTTTGCGTATTGGGCGCTCTTCCGCTTCCTCGCTCACTGACTCGCTGCGCTCGGTCGTTCGGCTGCGGCGAGCGGTATCAGCTCACTCAAAGGCGGTAATACGGTTATCCACAGAATCAGGGGATAACGCAGGAAAGAACATGTGAGCAAAAGGCCAGCAAAAGGCCAGGAACCGTAAAAAGGCCGCGTTGCTGGCGTTTTTCCATAGGCTCCGCCCCCCTGACGAGCATCACAAAAATCGACGCTCAAGTCAGAGGTGGCGAAACCCGACAGGACTATAAAGATACCAGGCGTTTCCCCCTGGAAGCTCCCTCGTGCGCTCTCCTGTTCCGACCCTGCCGCTTACCGGATACCTGTCCGCCTTTCTCCCTTCGGGAAGCGTGGCGCTTTCTCATAGCTCACGCTGTAGGTATCTCAGTTCGGTGTAGGTCGTTCGCTCCAAGCTGGGCTGTGTGCACGAACCCCCCGTTCAGCCCGACCGCTGCGCCTTATCCGGTAACTATCGTCTTGAGTCCAACCCGGTAAGACACGACTTATCGCCACTGGCAGCAGCCACTGGTAACAGGATTAGCAGAGCGAGGTATGTAGGCGGTGCTACAGAGTTCTTGAAGTGGTGGCCTAACTACGGCTACACTAGAAGAACAGTATTTGGTATCTGCGCTCTGCTGAAGCCAGTTACCTTCGGAAAAAGAGTTGGTAGCTCTTGATCCGGCAAACAAACCACCGCTGGTAGCGGTGGTTTTTTTGTTTGCAAGCAGCAGATTACGCGCAGAAAAAAAGGATCTCAAGAAGATCCTTTGATCTTTTCTACGGGGTCTGACGCTCAGTGGAACGAAAACTCACGTTAAGGGATTTTGGTCATGAGATTATCAAAAAGGATCTTCACCTAGATCCTTTTAAATTAAAAATGAAGTTTTAAATCAATCTAAAGTATATATGAGTAAACTTGGTCTGACAGTTACCAATGCTTAATCAGTGAGGCACCTATCTCAGCGATCTGTCTATTTCGTTCATCCATAGTTGCCTGACTCCCCGTCGTGTAGATAACTACGATACGGGAGGGCTTACCATCTGGCCCCAGTGCTGCAATGATACCGCGAGACCCACGCTCACCGGCTCCAGATTTATCAGCAATAAACCAGCCAGCCGGAAGGGCCGAGCGCAGAAGTGGTCCTGCAACTTTATCCGCCTCCATCCAGTCTATTAATTGTTGCCGGGAAGCTAGAGTAAGTAGTTCGCCAGTTAATAGTTTGCGCAACGTTGTTGCCATTGCTACAGGCATCGTGGTGTCACGCTCGTCGTTTGGTATGGCTTCATTCAGCTCCGGTTCCCAACGATCAAGGCGAGTTACATGATCCCCCATGTTGTGCAAAAAAGCGGTTAGCTCCTTCGGTCCTCCGATCGTTGTCAGAAGTAAGTTGGCCGCAGTGTTATCACTCATGGTTATGGCAGCACTGCATAATTCTCTTACTGTCATGCCATCCGTAAGATGCTTTTCTGTGACTGGTGAGTACTCAACCAAGTCATTCTGAGAATAGTGTATGCGGCGACCGAGTTGCTCTTGCCCGGCGTCAATACGGGATAATACCGCGCCACATAGCAGAACTTTAAAAGTGCTCATCATTGGAAAACGTTCTTCGGGGCGAAAACTCTCAAGGATCTTACCGCTGTTGAGATCCAGTTCGATGTAACCCACTCGTGCACCCAACTGATCTTCAGCATCTTTTACTTTCACCAGCGTTTCTGGGTGAGCAAAAACAGGAAGGCAAAATGCCGCAAAAAAGGGAATAAGGGCGACACGGAAATGTTGAATACTCATACTCTTCCTTTTTCAATATTATTGAAGCATTTATCAGGGTTATTGTCTCATGAGCGGATACATATTTGAATGTATTTAGAAAAATAAACAAATAGGGGTTCCGCGCACATTTCCCCGAAAAGTGCCACCTAAATTGTAAGCGTTAATATTTTGTTAAAATTCGCGTTAAATTTTTGTTAAATCAGCTCATTTTTTAACCAATAGGCCGAAATCGGCAAAATCCCTTATAAATCAAAAGAATAGACCGAGATAGGGTTGAGTGTTGTTCCAGTTTGGAACAAGAGTCCACTATTAAAGAACGTGGACTCCAACGTCAAAGGGCGAAAAACCGTCTATCAGGGCGATGGCCCACTACGTGAACCATCACCCTAATCAAGTTTTTTGGGGTCGAGGTGCCGTAAAGCACTAAATCGGAACCCTAAAGGGAGCCCCCGATTTAGAGCTTGACGGGGAAAGCCGGCGAACGTGGCGAGAAAGGAAGGGAAGAAAGCGAAAGGAGCGGGCGCTAGGGCGCTGGCAAGTGTAGCGGTCACGCTGCGCGTAACCACCACACCCGCCGCGCTTAATGCGCCGCTACAGGGCGCGTCCCATTCGCCATTCAGGCTGCGCAACTGTTGGGAAGGGCGATCGGTGCGGGCCTCTTCGCTATTACGCCAGCTGGCGAAAGGGGGATGTGCTGCAAGGCGATTAAGTTGGGTAACGCCAGGGTTTTCCCAGTCACGACGTTGTAAAACGACGGCCAGTGAATTGTAATACGACTCACTATAGGACGAAGACAAACAAACCATTATTATCATTAAAAGGCTCAGGAGAAACTTTAACAGTAATCAAAATGTCTGTTACAGTCAAGAGAATCATTGACAACACAGTCGTAGTTCCAAAACTTCCTGCAAATGAGGATCCAGTGGAATACCCGGCAGATTACTTCAGAAAATCAAAGGAGATTCCTCTTTACATCAATACTACAAAAAGTTTGTCAGATCTAAGAGGATATGTCTACCAAGGCCTCAAATCCGGAAATGTATCAATCATACATGTCAACAGCTACTTGTATGGAGCATTAAAGGACATCCGGGGTAAGTTGGATAAAGATTGGTCAAGTTTCGGAATAAACATCGGGAAAGCAGGGGATACAATCGGAATATTTGACCTTGTATCCTTGAAAGCCCTGGACGGCGTACTTCCAGATGGAGTATCGGATGCTTCCAGAACCAGCGCAGATGACAAATGGTTGCCTTTGTATCTACTTGGCTTATACAGAGTGGGCAGAACACAAATGCCTGAATACAGAAAAAAGCTCATGGATGGGCTGACAAATCAATGCAAAATGATCAATGAACAGTTTGAACCTCTTGTGCCAGAAGGTCGTGACATTTTTGATGTGTGGGGAAATGACAGTAATTACACAAAAATTGTCGCTGCAGTGGACATGTTCTTCCACATGTTCAAAAAACATGAATGTGCCTCGTTCAGATACGGAACTATTGTTTCCAGATTCAAAGATTGTGCTGCATTGGCAACATTTGGACACCTCTGCAAAATAACCGGAATGTCTACAGAAGATGTAACGACCTGGATCTTGAACCGAGAAGTTGCAGATGAAATGGTCCAAATGATGCTTCCAGGCCAAGAAATTGACAAGGCCGATTCATACATGCCTTATTTGATCGACTTTGGATTGTCTTCTAAGTCTCCATATTCTTCCGTCAAAAACCCTGCCTTCCACTTCTGGGGGCAATTGACAGCTCTTCTGCTCAGATCCACCAGAGCAAGGAATGCCCGACAGCCTGATGACATTGAGTATACATCTCTTACTACAGCAGGTTTGTTGTACGCTTATGCAGTAGGATCCTCTGCCGACTTGGCACAACAGTTTTGTGTTGGAGATAACAAATACACTCCAGATGATAGTACCGGAGGATTGACGACTAATGCACCGCCACAAGGCAGAGATGTGGTCGAATGGCTCGGATGGTTTGAAGATCAAAACAGAAAACCGACTCCTGATATGATGCAGTATGCGAAAAGAGCAGTCATGTCACTGCAAGGCCTAAGAGAGAAGACAATTGGCAAGTATGCTAAGTCAGAATTTGACAAATGACCCTATAATTCTCAGATCAC |
| F2 | 1,751 | TGACAAATGACCCTATAATTCTCAGATCACCTATTATATATTATGCTACATATGAAAAAAACTAACAGATATCATGGATAATCTCACAAAAGTTCGTGAGTATCTCAAGTCCTATTCTCGTCTGGATCAGGCGGTAGGAGAGATAGATGAGATCGAAGCACAACGAGCTGAAAAGTCCAATTATGAGTTGTTCCAAGAGGATGGAGTGGAAGAGCATACTAAGCCCTCTTATTTTCAGGCAGCAGATGATTCTGACACAGAATCTGAACCAGAAATTGAAGACAATCAAGGCTTGTATGCACCAGATCCAGAAGCTGAGCAAGTTGAAGGCTTTATACAGGGGCCTTTAGATGACTATGCAGATGAGGAAGTGGATGTTGTATTTACTTCGGACTGGAAACAGCCTGAGCTTGAATCTGACGAGCATGGAAAGACCTTACGGTTGACATCGCCAGAGGGTTTAAGTGGAGAGCAGAAATCCCAGTGGCTTTCGACGATTAAAGCAGTCGTGCAAAGTGCCAAATACTGGAATCTGGCAGAGTGCACATTTGAAGCATCGGGAGAAGGGGTCATTATGAAGGAGCGCCAGATAACTCCGGATGTATATAAGGTCACTCCAGTGATGAACACACATCCGTCCCAATCAGAAGCAGTATCAGATGTTTGGTCTCTCTCAAAGACATCCATGACTTTCCAACCCAAGAAAGCAAGTCTTCAGCCTCTCACCATATCCTTGGATGAATTGTTCTCATCTAGAGGAGAGTTCATCTCTGTCGGAGGTGACGGACGAATGTCTCATAAAGAGGCCATCCTGCTCGGCCTGAGATACAAAAAGTTGTACAATCAGGCGAGAGTCAAATATTCTCTGTAGACTATGAAAAAAAGTAACAGATATCACGATCTAAGTGTTATCCCAATCCATTCATCATGAGTTCCTTAAAGAAGATTCTCGGTCTGAAGGGGAAAGGTAAGAAATCTAAGAAATTAGGGATCGCACCACCCCCTTATGAAGAGGACACTAGCATGGAGTATGCTCCGAGCGCTCCAATTGACAAATCCTATTTTGGAGTTGACGAGATGGACACCTATGATCCGAATCAATTAAGATATGAGAAATTCTTCTTTACAGTGAAAATGACGGTTAGATCTAATCGTCCGTTCAGAACATACTCAGATGTGGCAGCCGCTGTATCCCATTGGGATCACATGTACATCGGAATGGCAGGGAAACGTCCCTTCTACAAAATCTTGGCTTTTTTGGGTTCTTCTAATCTAAAGGCCACTCCAGCGGTATTGGCAGATCAAGGTCAACCAGAGTATCACGCTCACTGCGAAGGCAGGGCTTATTTGCCACATAGGATGGGGAAGACCCCTCCCATGCTCAATGTACCAGAGCACTTCAGAAGACCATTCAATATAGGTCTTTACAAGGGAACGATTGAGCTCACAATGACCATCTACGATGATGAGTCACTGGAAGCAGCTCCTATGATCTGGGATCATTTCAATTCTTCCAAATTTTCTGATTTCAGAGAGAAGGCCTTAATGTTTGGCCTGATTGTCGAGAAAAAGGCATCTGGAGCGTGGGTCCTGGACTCTATCGGCCACTTCAAATGAGCTAGTCTAACTTCTAGCTTCTGAACAATCCCCGGTTTACTCAGTCTCCCCTAATTCCAGCCTCTCGAACAACTAATATCCTGTCTTTTCTATCCCTATGAAAAAAACTAACAGAGATCGATCTGTTTACGCGT |
| F3 | 1,709 | AAACTAACAGAGATCGATCTGTTTACGCGTCACTATGAAGTGCCTTTTGTACTTAGCCTTTTTATTCATTGGGGTGAATTGCAAGTTCACCATAGTTTTTCCACACAACCAAAAAGGAAACTGGAAAAATGTTCCTTCTAATTACCATTATTGCCCGTCAAGCTCAGATTTAAATTGGCATAATGACTTAATAGGCACAGCCTTACAAGTCAAAATGCCCAAGAGTCACAAGGCTATTCAAGCAGACGGTTGGATGTGTCATGCTTCCAAATGGGTCACTACTTGTGATTTCCGCTGGTATGGACCGAAGTATATAACACATTCCATCCGATCCTTCACTCCATCTGTAGAACAATGCAAGGAAAGCATTGAACAAACGAAACAAGGAACTTGGCTGAATCCAGGCTTCCCTCCTCAAAGTTGTGGATATGCAACTGTGACGGATGCCGAAGCAGTGATTGTCCAGGTGACTCCTCACCATGTGCTGGTTGATGAATACACAGGAGAATGGGTTGATTCACAGTTCATCAACGGAAAATGCAGCAATTACATATGCCCCACTGTCCATAACTCTACAACCTGGCATTCTGACTATAAGGTCAAAGGGCTATGTGATTCTAACCTCATTTCCATGGACATCACCTTCTTCTCAGAGGACGGAGAGCTATCATCCCTGGGAAAGGAGGGCACAGGGTTCAGAAGTAACTACTTTGCTTATGAAACTGGAGGCAAGGCCTGCAAAATGCAATACTGCAAGCATTGGGGAGTCAGACTCCCATCAGGTGTCTGGTTCGAGATGGCTGATAAGGATCTCTTTGCTGCAGCCAGATTCCCTGAATGCCCAGAAGGGTCAAGTATCTCTGCTCCATCTCAGACCTCAGTGGATGTAAGTCTAATTCAGGACGTTGAGAGGATCTTGGATTATTCCCTCTGCCAAGAAACCTGGAGCAAAATCAGAGCGGGTCTTCCAATCTCTCCAGTGGATCTCAGCTATCTTGCTCCTAAAAACCCAGGAACCGGTCCTGCTTTCACCATAATCAATGGTACCCTAAAATACTTTGAGACCAGATACATCAGAGTCGATATTGCTGCTCCAATCCTCTCAAGAATGGTCGGAATGATCAGTGGAACTACCACAGAAAGGGAACTGTGGGATGACTGGGCACCATATGAAGACGTGGAAATTGGACCCAATGGAGTTCTGAGGACCAGTTCAGGATATAAGTTTCCTTTATACATGATTGGACATGGTATGTTGGACTCCGATCTTCATCTTAGCTCAAAGGCTCAGGTGTTCGAACATCCTCACATTCAAGACGCTGCTTCGCAACTTCCTGATGATGAGAGTTTATTTTTTGGTGATACTGGGCTATCCAAAAATCCAATCGAGCTTGTAGAAGGTTGGTTCAGTAGTTGGAAAAGCTCTATTGCCTCTTTTTTCTTTATCATAGGGTTAATCATTGGACTATTCTTGGTTCTCCGAGTTGGTATCCATCTTTGCATTAAATTAAAGCACACCAAGAAAAGACAGATTTATACAGACATAGAGATGAACCGACTTGGAAAGTAACTCAAATCCTGCTAGCCAGATTCTTCATGTTTGGACCAAATCAACTTGTGATACCATGCTCAAAGAGGCCTCAATTATATTTGAGTTTTTAATTTTTATGAAAAAAACTAACAGCAATCATGGAAGTCCACGATTTTGA |
| F4 | 6,439 | AACAGCAATCATGGAAGTCCACGATTTTGAGACCGACGAGTTCAATGATTTCAATGAAGATGACTATGCCACAAGAGAATTCCTGAATCCCGATGAGCGCATGACGTACTTGAATCATGCTGATTACAACCTGAATTCTCCTCTAATTAGTGATGATATTGACAATTTAATCAGGAAATTCAATTCTCTTCCAATTCCCTCGATGTGGGATAGTAAGAACTGGGATGGAGTTCTTGAGATGTTAACATCATGTCAAGCCAATCCCATCTCAACATCTCAGATGCATAAATGGATGGGAAGTTGGTTAATGTCTGATAATCATGATGCCAGTCAAGGGTATAGTTTTTTACATGAAGTGGACAAAGAGGCAGAAATAACATTTGACGTGGTGGAGACCTTCATCCGCGGCTGGGGCAACAAACCAATTGAATACATCAAAAAGGAAAGATGGACTGACTCATTCAAAATTCTCGCTTATTTGTGTCAAAAGTTTTTGGACTTACACAAGTTGACATTAATCTTAAATGCTGTCTCTGAGGTGGAATTGCTCAACTTGGCGAGGACTTTCAAAGGCAAAGTCAGAAGAAGTTCTCATGGAACGAACATATGCAGGATTAGGGTTCCCAGCTTGGGTCCTACTTTTATTTCAGAAGGATGGGCTTACTTCAAGAAACTTGATATTCTAATGGACCGAAACTTTCTGTTAATGGTCAAAGATGTGATTATAGGGAGGATGCAAACGGTGCTATCCATGGTATGTAGAATAGACAACCTGTTCTCAGAGCAAGACATCTTCTCCCTTCTAAATATCTACAGAATTGGAGATAAAATTGTGGAGAGGCAGGGAAATTTTTCTTATGACTTGATTAAAATGGTGGAACCGATATGCAACTTGAAGCTGATGAAATTAGCAAGAGAATCAAGGCCTTTAGTCCCACAATTCCCTCATTTTGAAAATCATATCAAGACTTCTGTTGATGAAGGGGCAAAAATTGACCGAGGTATAAGATTCCTCCATGATCAGATAATGAGTGTGAAAACAGTGGATCTCACACTGGTGATTTATGGATCGTTCAGACATTGGGGTCATCCTTTTATAGATTATTACACTGGACTAGAAAAATTACATTCCCAAGTAACCATGAAGAAAGATATTGATGTGTCATATGCAAAAGCACTTGCAAGTGATTTAGCTCGGATTGTTCTATTTCAACAGTTCAATGATCATAAAAAGTGGTTCGTGAATGGAGACTTGCTCCCTCATGATCATCCCTTTAAAAGTCATGTTAAAGAAAATACATGGCCCACAGCTGCTCAAGTTCAAGATTTTGGAGATAAATGGCATGAACTTCCGCTGATTAAATGTTTTGAAATACCCGACTTACTAGACCCATCGATAATATACTCTGACAAAAGTCATTCAATGAATAGGTCAGAGGTGTTGAAACATGTCCGAATGAATCCGAACACTCCTATCCCTAGTAAAAAGGTGTTGCAGACTATGTTGGACACAAAGGCTACCAATTGGAAAGAATTTCTTAAAGAGATTGATGAGAAGGGCTTAGATGATGATGATCTAATTATTGGTCTTAAAGGAAAGGAGAGGGAACTGAAGTTGGCAGGTAGATTTTTCTCCCTAATGTCTTGGAAATTGCGAGAATACTTTGTAATTACCGAATATTTGATAAAGACTCATTTCGTCCCTATGTTTAAAGGCCTGACAATGGCGGACGATCTAACTGCAGTCATTAAAAAGATGTTAGATTCCTCATCCGGCCAAGGATTGAAGTCATATGAGGCAATTTGCATAGCCAATCACATTGATTACGAAAAATGGAATAACCACCAAAGGAAGTTATCAAACGGCCCAGTGTTCCGAGTTATGGGCCAGTTCTTAGGTTATCCATCCTTAATCGAGAGAACTCATGAATTTTTTGAGAAAAGTCTTATATACTACAATGGAAGACCAGACTTGATGCGTGTTCACAACAACACACTGATCAATTCAACCTCCCAACGAGTTTGTTGGCAAGGACAAGAGGGTGGACTGGAAGGTCTACGGCAAAAAGGATGGAGTATCCTCAATCTACTGGTTATTCAAAGAGAGGCTAAAATCAGAAACACTGCTGTCAAAGTCTTGGCACAAGGTGATAATCAAGTTATTTGCACACAGTATAAAACGAAGAAATCGAGAAACGTTGTAGAATTACAGGGTGCTCTCAATCAAATGGTTTCTAATAATGAGAAAATTATGACTGCAATCAAAATAGGGACAGGGAAGTTAGGACTTTTGATAAATGACGATGAGACTATGCAATCTGCAGATTACTTGAATTATGGAAAAATACCGATTTTCCGTGGAGTGATTAGAGGGTTAGAGACCAAGAGATGGTCACGAGTGACTTGTGTCACCAATGACCAAATACCCACTTGTGCTAATATAATGAGCTCAGTTTCCACAAATGCTCTCACCGTAGCTCATTTTGCTGAGAACCCAATCAATGCCATGATACAGTACAATTATTTTGGGACATTTGCTAGACTCTTGTTGATGATGCATGATCCTGCTCTTCGTCAATCATTGTATGAAGTTCAAGATAAGATACCGGGCTTGCACAGTTCTACTTTCAAATACGCCATGTTGTATTTGGACCCTTCCATTGGAGGAGTGTCGGGCATGTCTTTGTCCAGGTTTTTGATTAGAGCCTTCCCAGATCCCGTAACAGAAAGTCTCTCATTCTGGAGATTCATCCATGTACATGCTCGAAGTGAGCATCTGAAGGAGATGAGTGCAGTATTTGGAAACCCCGAGATAGCCAAGTTTCGAATAACTCACATAGACAAGCTAGTAGAAGATCCAACCTCTCTGAACATCGCTATGGGAATGAGTCCAGCGAACTTGTTAAAGACTGAGGTTAAAAAATGCTTAATCGAATCAAGACAAACCATCAGGAACCAGGTGATTAAGGATGCAACCATATATTTGTATCATGAAGAGGATCGGCTCAGAAGTTTCTTATGGTCAATAAATCCTCTGTTCCCTAGATTTTTAAGTGAATTCAAATCAGGCACTTTTTTGGGAGTCGCAGACGGGCTCATCAGTCTATTTCAAAATTCTCGTACTATTCGGAACTCCTTTAAGAAAAAGTATCATAGGGAATTGGATGATTTGATTGTGAGGAGTGAGGTATCCTCTTTGACACATTTAGGGAAACTTCATTTGAGAAGGGGATCATGTAAAATGTGGACATGTTCAGCTACTCATGCTGACACATTAAGATACAAATCCTGGGGCCGTACAGTTATTGGGACAACTGTACCCCATCCATTAGAAATGTTGGGTCCACAACATCGAAAAGAGACTCCTTGTGCACCATGTAACACATCAGGGTTCAATTATGTTTCTGTGCATTGTCCAGACGGGATCCATGACGTCTTTAGTTCACGGGGACCATTGCCTGCTTATCTAGGGTCTAAAACATCTGAATCTACATCTATTTTGCAGCCTTGGGAAAGGGAAAGCAAAGTCCCACTGATTAAAAGAGCTACACGTCTTAGAGATGCTATCTCTTGGTTTGTTGAACCCGACTCTAAACTAGCAATGACTATACTTTCTAACATCCACTCTTTAACAGGCGAAGAATGGACCAAAAGGCAGCATGGGTTCAAAAGAACAGGGTCTGCCCTTCATAGGTTTTCGACATCTCGGATGAGCCATGGTGGGTTCGCATCTCAGAGCACTGCAGCATTGACCAGGTTGATGGCAACTACAGACACCATGAGGGATCTGGGAGATCAGAATTTCGACTTTTTATTCCAAGCAACGTTGCTCTATGCTCAAATTACCACCACTGTTGCAAGAGACGGATGGATCACCAGTTGTACAGATCATTATCATATTGCCTGTAAGTCCTGTTTGAGACCCATAGAAGAGATCACCCTGGACTCAAGTATGGACTACACGCCCCCAGATGTATCCCATGTGCTGAAGACATGGAGGAATGGGGAAGGTTCGTGGGGACAAGAGATAAAACAGATCTATCCTTTAGAAGGGAATTGGAAGAATTTAGCACCTGCTGAGCAATCCTATCAAGTCGGCAGATGTATAGGTTTTCTATATGGAGACTTGGCGTATAGAAAATCTACTCATGCCGAGGACAGTTCTCTATTTCCTCTATCTATACAAGGTCGTATTAGAGGTCGAGGTTTCTTAAAAGGGTTGCTAGACGGATTAATGAGAGCAAGTTGCTGCCAAGTAATACACCGGAGAAGTCTGGCTCATTTGAAGAGGCCGGCCAACGCAGTGTACGGAGGTTTGATTTACTTGATTGATAAATTGAGTGTATCACCTCCATTCCTTTCTCTTACTAGATCAGGACCTATTAGAGACGAATTAGAAACGATTCCCCACAAGATCCCAACCTCCTATCCGACAAGCAACCGTGATATGGGGGTGATTGTCAGAAATTACTTCAAATACCAATGCCGTCTAATTGAAAAGGGAAAATACAGATCACATTATTCACAATTATGGTTATTCTCAGATGTCTTATCCATAGACTTCATTGGACCATTCTCTATTTCCACCACCCTCTTGCAAATCCTATACAAGCCATTTTTATCTGGGAAAGATAAGAATGAGTTGAGAGAGCTGGCAAATCTTTCTTCATTGCTAAGATCAGGAGAGGGGTGGGAAGACATACATGTGAAATTCTTCACCAAGGACATATTATTGTGTCCAGAGGAAATCAGACATGCTTGCAAGTTCGGGATTGCTAAGGATAATAATAAAGACATGAGCTATCCCCCTTGGGGAAGGGAATCCAGAGGGACAATTACAACAATCCCTGTTTATTATACGACCACCCCTTACCCAAAGATGCTAGAGATGCCTCCAAGAATCCAAAATCCCCTGCTGTCCGGAATCAGGTTGGGCCAATTACCAACTGGCGCTCATTATAAAATTCGGAGTATATTACATGGAATGGGAATCCATTACAGGGACTTCTTGAGTTGTGGAGACGGCTCCGGAGGGATGACTGCTGCATTACTACGAGAAAATGTGCATAGCAGAGGAATATTCAATAGTCTGTTAGAATTATCAGGGTCAGTCATGCGAGGCGCCTCTCCTGAGCCCCCCAGTGCCCTAGAAACTTTAGGAGGAGATAAATCGAGATGTGTAAATGGTGAAACATGTTGGGAATATCCATCTGACTTATGTGACCCAAGGACTTGGGACTATTTCCTCCGACTCAAAGCAGGCTTGGGGCTTCAAATTGATTTAATTGTAATGGATATGGAAGTTCGGGATTCTTCTACTAGCCTGAAAATTGAGACGAATGTTAGAAATTATGTGCACCGGATTTTGGATGAGCAAGGAGTTTTAATCTACAAGACTTATGGAACATATATTTGTGAGAGCGAAAAGAATGCAGTAACAATCCTTGGTCCCATGTTCAAGACGGTCGACTTAGTTCAAACAGAATTTAGTAGTTCTCAAACGTCTGAAGTATATATGGTATGTAAAGGTTTGAAGAAATTAATCGATGAACCCAATCCCGATTGGTCTTCCATCAATGAATCCTGGAAAAACCTGTACGCATTCCAGTCATCAGAACAGGAATTTGCCAGAGCAAAGAAGGTTAGTACATACTTTACCTTGACAGGTATTCCCTCCCAATTCATTCCTGATCCTTTTGTAAACATTGAGACTATGCTACAAATATTCGGAGTACCCACGGGTGTGTCTCATGCGGCTGCCTTAAAATCATCTGATAGACCTGCAGATTTATTGACCATTAGCCTTTTTTATATGGCGATTATATCGTATTATAACATCAATCATATCAGAGTAGGACCGATACCTCCGAACCCCCCATCAGATGGAATTGCACAAAATGTGGGGATCGCTATAACTGGTATAAGCTTTTGGCTGAGTTTGATGGAGAAAGACATTCCACTATATCAACAGTGTTTAGCAGTTATCCAGCAATCATTCCCGATTAGGTGGGAGGCTGTTTCAGTAAAAGGAGGATACAAGCAGAAGTGGAGTACTAGAGGTGATGGGCTCCCAAAAGATACCCGAATTTCAGACTCCTTGGCCCCAATCGGGAACTGGATCAGATCTCTGGAATTGGTCCGAAACCAAGTTCGTCTAAATCCATTCAATGAGATCTTGTTCAATCAGCTATGTCGTACAGTGGATAATCATTTGAAATGGTCAAATTTGCGAAGAAACACAGGAATGATTGAATGGATCAATAGACGAATTTCAAAAGAAGACCGGTCTATACTGATGTTGAAGAGTGACCTACACGAGGAAAACTCTTGGAGAGATTAAAAAATCATGAGGAGACTCCAAACTTTAAGTATGAAAAAAACTTTGATCCTTAAGACCCTCTTGTGGTTTTTATTTTTTATCTGGTTTTGTGGTCTTCGT |
